# Data-driven yield projections suggest large opportunities to improve Europe’s soybean self-sufficiency under climate change

**DOI:** 10.1101/2020.10.08.331496

**Authors:** Nicolas Guilpart, Toshichika Iizumi, David Makowski

## Abstract

Currently, demand for soybean in Europe is mostly fulfilled by imports. However, soybean-growing areas across Europe have been rapidly increasing in response to a rising demand for locally-produced, non-GM soybean in recent years. This raises questions about the suitability of European agro-climatic conditions for soybean production. We used data-driven relationships between climate and soybean yield derived from machine-learning techniques to make yield projections under current and future climate with moderate (RCP 4.5) to intense (RCP 8.5) warming, up to the 2050s and 2090s time horizons. Results suggest that a self-sufficiency level of 50% (100%) would be achievable in Europe under historical and future climate if 4-5% (9-12%) of the current European cropland is dedicated to soybean production. The associated increase in soybean area in Europe would bring environmental benefits, with a potential decrease of nitrogen fertilizer use in Europe by 5-8% (13-18%) and a possible reduction of deforestation in biodiversity hotspots in South America. However, it would also lead to an important reduction in the production of other cultivated species in Europe (e.g. cereals) and a potential increase in the use of irrigation water.

## Introduction

The satisfaction of European^*^ soybean demand is highly dependent on imports. Currently, Europe imports about 58 Mt yr^-1^ of soybean which accounts for nearly 90% of the domestic consumption^1^ (average over 2009-2013; Table S1). This large share of soybean imports in Europe takes it roots in the post-World War II international trade agreements between Europe and the USA that allowed tax-free entry of protein imports into Europe^2^. Price support for cereals cultivated within the European Economic Community led to a strong growth of cereal production in Europe at the expense of grain legumes^3^. This political context explains why the extent of legume production area has been limited in Europe, despite the increasing demand. Only 1.7% of European cropland area was used for soybean production in 2016^1^. However, it is well-documented that legume (including soybean) production and consumption have numerous benefits. First, it increases yield of the subsequent crop and reduces occurrence of weeds and pathogens ^4–6^ (agronomic benefit). Second, it reduces use of nitrogen (N) fertilizer due to symbiotic N fixation and associated reductions in greenhouse gases emissions and energy use^5,7^ (environmental benefit). Lastly, legume consumption contributes to reducing risks associated with chronic diseases, such as cardiovascular diseases, diabetes, cancer, obesity and gut health^8^ (human health benefit, see ^9–11^ for soybean). On the other hand, increases in legume-producing areas may lead to side effects. For example, in comparison to cereals, soybean production less contributes to soil carbon sequestration^12,13^. It may also lead to an increased reliance on pesticides (e.g. pea^14^) or irrigation (e.g. soybean^15^). Despite these side effects, the overall benefits of increasing the share of legume crops in European agricultural systems are still expected to be positive^8^.

Among commonly cultivated grain legumes, soybean stands out as the crop species experiencing the fastest expansion rate in Europe with an increase of more than four-fold from 1.2 Mha in 2004 to 5 Mha in 2016 (Figure S1) in response to a rising demand for locally-produced, non-GM soybean^16–18^. This expansion is expected to continue in the next decade but at a slower pace^19^. In this context, a Europe-wide assessment on the agro-climatic suitability of soybean production areas under current and future climate is of strategic importance. Building on two recently published global datasets including historical soybean yield ^20,21^ and retrospective meteorological forcing^22^, we developed data-driven relationships between climate and soybean yield to estimate soybean suitable areas over Europe. Several machine learning algorithms were trained and tested at the global scale (Random Forest, Artificial Neural Networks, Generalized Additive Model, and Multiple Linear Regression) to predict soybean yield as a function of monthly climate inputs (solar radiation, minimum and maximum temperature, rainfall, and vapour pressure) calculated over the growing season (April to October). A large share of the training data was taken from major soybean-producing countries (Argentina, Brazil, Canada, China, India, Italy and the United States), and zero-yield data points were randomly sampled in climate zones known to be unsuitable for soybean production (e.g. deserts and arctic areas) and added to the dataset so that they represented about 20% of the final dataset. The most accurate algorithm was selected after running a cross-validation procedure assessing model transferability in time and space^23,24^. The selected algorithm (Random Forest) was then run for the entire Europe to assess potential distribution of soybean suitable area in rainfed conditions under current and future climate. Projections of soybean suitability in Europe were performed using 16 climate change scenarios consisting of bias-corrected data produced by eight Global Climate Models of the Coupled Model Intercomparison phase 5 (CMIP5)^25^ and two Representative Concentration Pathways (RCPs; 4.5 and 8.5 W m^-2^) ^26^ in the 2050s and the 2090s. The projections assume a growing season from April to October and no irrigation, although soybean is often irrigated in Europe^15^. The no irrigation assumption prevents from making any hypothesis about available water for irrigation, which is a complex issue especially under climate change^27,28^. We therefore acknowledge that our yield projections are probably a bit conservative from that point of view. Day length, soil type and atmospheric CO2 concentration are other factors not accounted for in our model, justification of these choices and implications for the results are discussed hereafter.

## Results

### Model fitting and selection

Among algorithms tested in this study, Random Forest appears to be the most accurate in terms of root-mean-squared error of prediction (RMSEP) and Nash–Sutcliffe model efficiency coefficient (MEF) (Table 1, Figure S2). It achieves the lowest prediction error (RMSEP = 0.35 t ha ^-1^) and the highest efficiency (MEF = 0.93), as estimated with an unstratified cross validation procedure. It also displays the best transferability in time (RMSEP = 0.45 t ha^-1^ when applied to years different from those used for training) and space (RMSEP = 0.43 t ha^-1^ when applied in locations distant by 500 km – or 5 grid-cells – from those used for training). Our results reveal that transferability in space decreases with increasing distance between training and test datasets for all models, with a threshold of 1000 km above which the performance of the selected algorithm deteriorates markedly (Table 1).

**Table 1.**
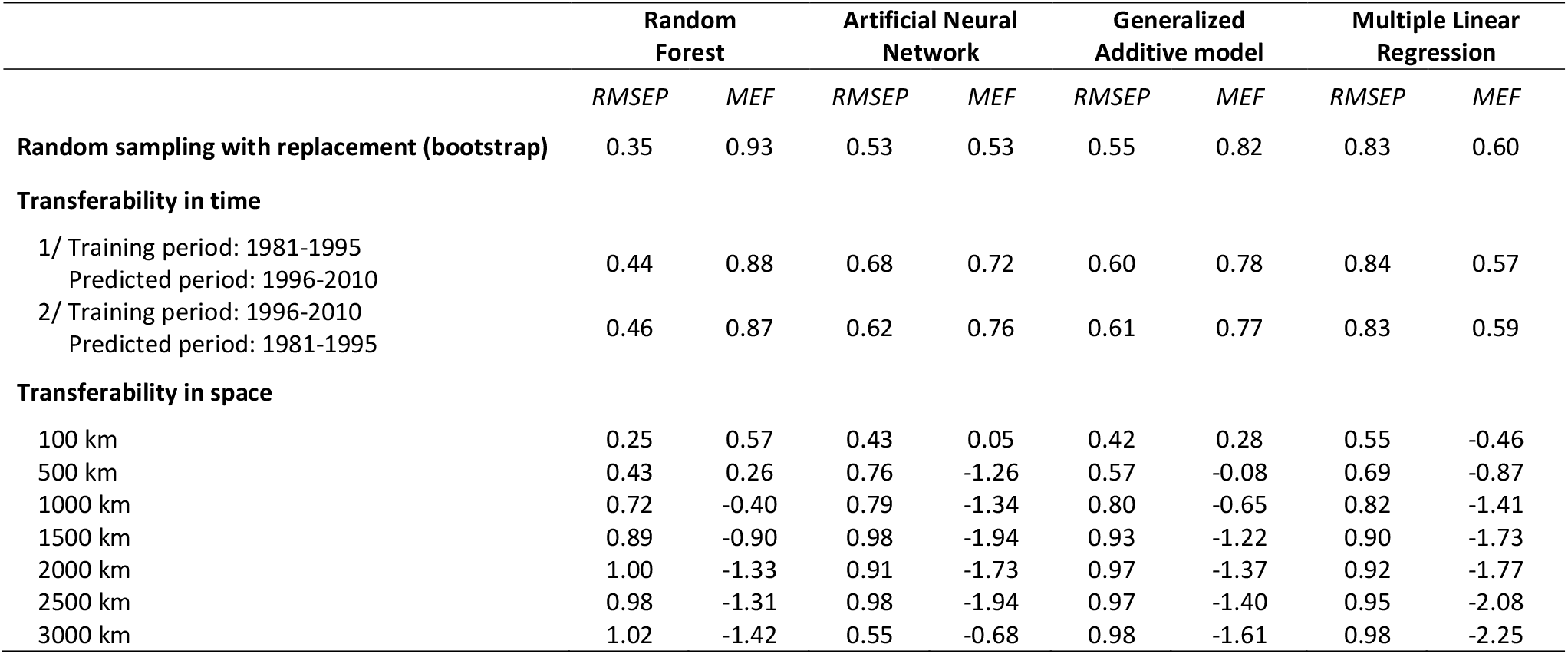
Performances assessment of the tested machine learning algorithms. Soybean yield was expressed as a function 5 climate variables (daily minimum and maximum temperatures, rainfall, solar radiation, and vapour pressure) calculated monthly for a 7-month growing season, plus the irrigated fraction in a grid-cell, making a total of 36 predictors. The whole dataset contains 30,337 yield observations from 1981 to 2010. The model predictive ability is first assessed using 25 out-of-bag samples generated by bootstrap. Then, transferability in time is assessed by fitting the algorithms on a period of time different than the predicted period, and transferability in space is assessed by ensuring a minimum spatial distance between training and test datasets, with seven minimal distances ranging from 100 km to 3000 km considered successively. RMSEP: root mean square error of prediction (t ha^-1^). RMSEP values for transferability in space are the median over 70 RMSEP values (10 grid-cells * 7 countries). MEF: Nash–Sutcliffe model efficiency. An efficiency of 1 corresponds to a perfect match of modeled to observed data, an efficiency of 0 indicates that model predictions are as accurate as the mean of observed data, whereas an efficiency lower than zero occurs when the observed mean is a better predictor than the model.

|  | Random Forest |  | Artificial Neural Network |  | Generalized Additive model |  | Multiple Linear Regression |  |
| --- | --- | --- | --- | --- | --- | --- | --- | --- |
|  | *RMSEP* | *MEF* | *RMSEP* | *MEF* | *RMSEP* | *MEF* | *RMSEP* | *MEF* |
| **Random sampling with replacement (bootstrap)** | 0.35 | 0.93 | 0.53 | 0.53 | 0.55 | 0.82 | 0.83 | 0.60 |
| **Transferability in time** |  |  |  |  |  |  |  |  |
| 1/ Training period: 1981-1995<br>Predicted period: 1996-2010 | 0.44 | 0.88 | 0.68 | 0.72 | 0.60 | 0.78 | 0.84 | 0.57 |
| 2/ Training period: 1996-2010<br>Predicted period: 1981-1995 | 0.46 | 0.87 | 0.62 | 0.76 | 0.61 | 0.77 | 0.83 | 0.59 |
| **Transferability in space** |  |  |  |  |  |  |  |  |
| 100 km | 0.25 | 0.57 | 0.43 | 0.05 | 0.42 | 0.28 | 0.55 | -0.46 |
| 500 km | 0.43 | 0.26 | 0.76 | -1.26 | 0.57 | -0.08 | 0.69 | -0.87 |
| 1000 km | 0.72 | -0.40 | 0.79 | -1.34 | 0.80 | -0.65 | 0.82 | -1.41 |
| 1500 km | 0.89 | -0.90 | 0.98 | -1.94 | 0.93 | -1.22 | 0.90 | -1.73 |
| 2000 km | 1.00 | -1.33 | 0.91 | -1.73 | 0.97 | -1.37 | 0.92 | -1.77 |
| 2500 km | 0.98 | -1.31 | 0.98 | -1.94 | 0.97 | -1.40 | 0.95 | -2.08 |
| 3000 km | 1.02 | -1.42 | 0.55 | -0.68 | 0.98 | -1.61 | 0.98 | -2.25 |

### Projections of soybean yield in Europe

The projections of the Random Forest algorithm – which assume no irrigation and a fixed growing period from April to October – suggest high suitability for soybean under historical and future climate (Figure 1). Under historical climate (Figure 1A), about 100 Mha show projected yield equal or higher than 2 t ha^-1^ (Figure 2B), while in 2016 the soybean production area in Europe was only 5 Mha with 2 t ha^-1^ of average yield^1^. Therefore, soybean suitable area appears to be much larger than current harvested area in Europe, which suggests that soybean production is not limited by climate conditions. Our projections indicate an overall positive effect of climate change on soybean yield, with a projected increase of median soybean yield from 1.2 t ha^-1^ under historical climate to 1.6 t ha^-1^ (2050s – RCP 4.5) and 1.8 t ha^-1^ (2090s – RCP 8.5), even without effects of elevated CO2 concentration (Figure 2A). Importantly, the increase in the extent of low-yielding suitable areas (+30% to +45% for areas with projected yield ? 1.5 t ha^-1^ relative to historical climate) was associated with a decrease in the extent of high-yielding suitable areas (−60% to −100% for areas with projected yield ≥ 2.5 t ha^-1^ relative to historical climate) (Figure 2B, Table S2). The decrease in medium-yielding suitable areas was substantial under RCP 8.5 (−20% to −54% relative to historical climate) compared to RCP 4.5 (−2% to −10%). These changes reflected losses in the South (e.g. Spain, Italy) and gains in the North and the East (e.g. Russia, Ukraine, Poland, and Belorussia). The northward and eastward shifts of higher-yielding suitable area and the decrease of suitable area in the South of Europe would become noticeable by the middle of this century (Figure 1B,D) and further intensify by the end of this century, in particular under RCP 8.5 (Figure 1C,E). We highlight that these projections do not involve any extrapolation of the model beyond the range of training data (historical growing season climate). Indeed, only 0.03% of the data samples in the future climate scenarios fall out of the range of training climate data (Figure S3, Table S3). Moreover, current available evidence from farmer’s fields and on-station field experiments (Figure S4-A) as well as from available estimates of current soybean harvested area in Europe (Figure S4-B) confirms that soybean can be grown at high latitude in Europe of 55°N to 57.5°N (corresponding to the northern part of Latvia).

**Figure 1.**
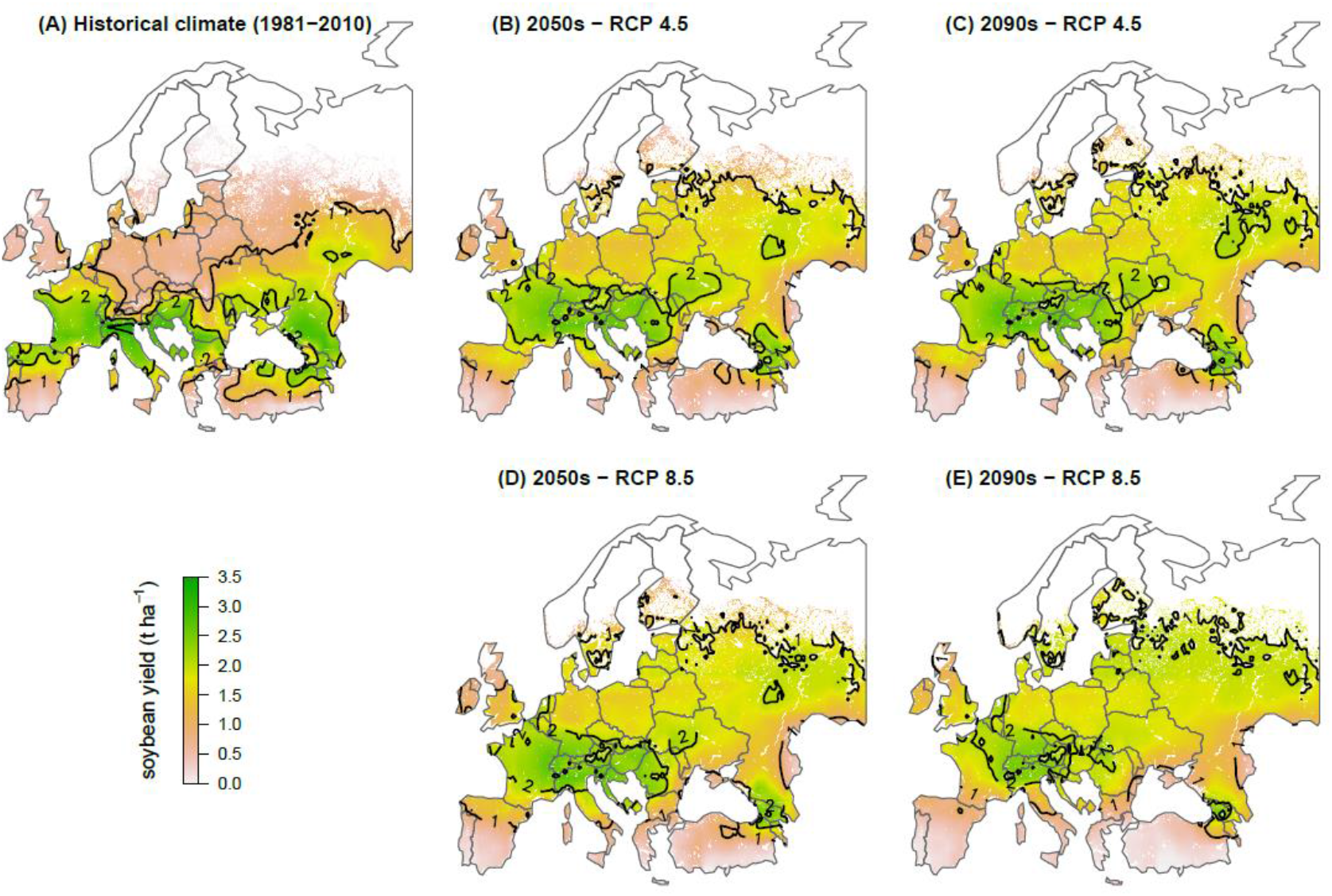
Projected soybean yield in Europe under historical and future climate. Projected soybean yield (A) under historical climate (1981-2010), (B) by mid-century (2050-2059) under RCP 4.5, (C) by the end of the century (2090-2099) under RCP 4.5, (D) by mid-century (2050-2059) under RCP 8.5, (E) by the end of the century (2090-2099) under RCP 8.5. Maps show median projected yield using a Random Forest algorithm run with the GRASP dataset^22^ for historical climate (1981-2010), and over the eight Global Circulation Models^25^ considered in this study for future climate scenarios. Projections are shown only on agricultural area (cropland plus pasture), in the year 2000^31^.

**Figure 2.**
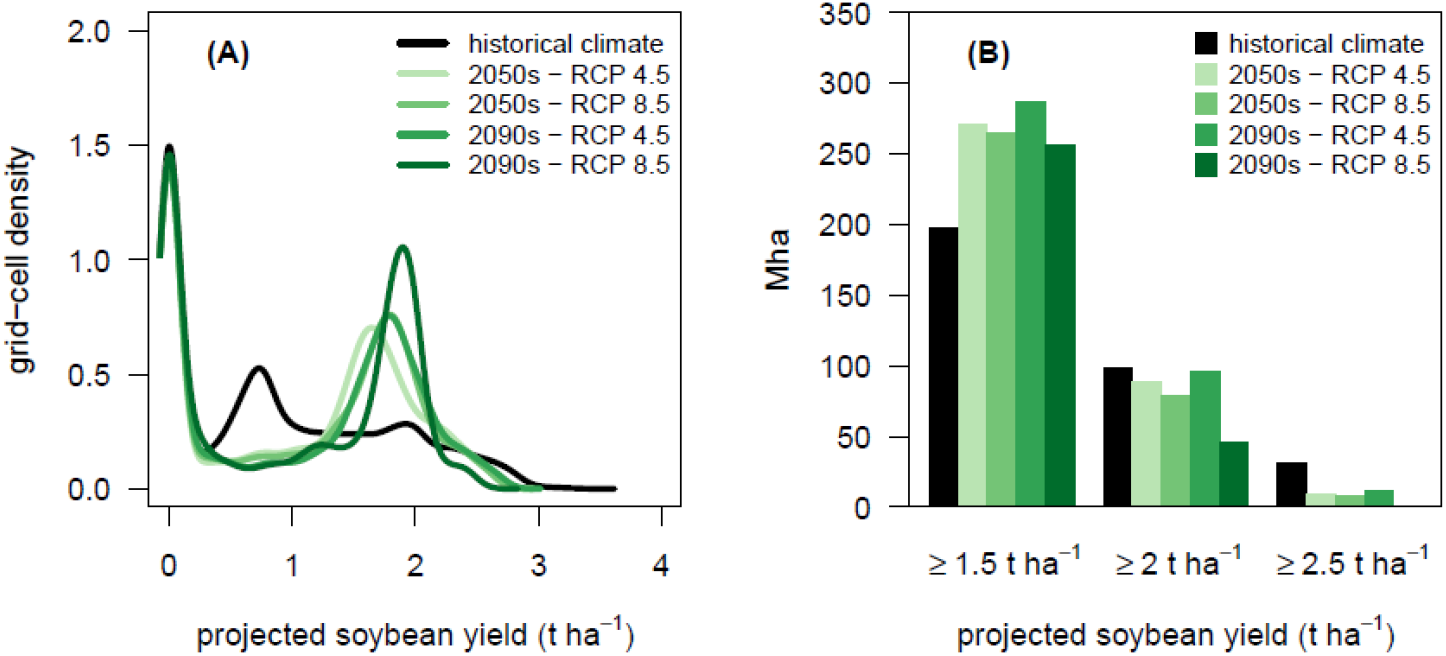
Effect of climate change on projected soybean yield in Europe. (A) Probability density functions of projected soybean yield under historical climate and different future climate scenarios. (B) Extent of European agricultural area for which projected soybean yield is higher or equal to a given yield threshold. Soybean yield projections were performed with a Random Forest algorithm. Projections for historical climate used the GRASP meteorological dataset^22^ from 1981 to 2010. For future climate, eight Global Circulation Models^25^ were used and median projected yield over the height model was calculated.

### Climate drivers of projected yield changes

To identify climate drivers of projected shifts in soybean suitability, we performed a linear discriminant analysis (LDA) to find out combinations of climate variables that best discriminate three types of yield response – yield decrease (by at least −0.3 t ha^-1^), yield increase (by at least +0.3 t ha^-1^), and a marginal change (projected yield change between −0.3 and +0.3 t ha^-1^) – when comparing RCP 4.5 in the 2050s to historical climate (Figure S5). The +/- 0.3 t ha^-1^ threshold was chosen to be higher than the observed interannual variability of soybean yield in Europe, which is 0.2 t ha^-1^ (standard deviation) over the 2000-2014 time period^1^. We also performed the same analysis with RCP 8.5 in the 2050s, but we present results for RCP 4.5 because conclusions are similar. The LDA showed an overall accuracy of 89%, and was able to discriminate between grid-cells experiencing yield increase or yield decrease (Figure 3A, Table S5). This analysis reveals the key role of temperature (both minimum and maximum) in driving projected yield changes. Indeed, climate variables showing the highest contributions to the first two linear discriminants are mostly temperature variables (Figure 3B-C, Figure S6). Our results suggest that projected yield decrease in the South of Europe is mainly associated with detrimental warming effects during the reproductive period in warmer regions. Maximum temperatures in months 4 and 5 of the growing season (i.e. July-August) reach 31.3°C and 30.9°C, respectively, under RCP 4.5 in the 2050s (Table 2), which exceeds the optimum of 28.5-30°C for pollen germination (see Table S7 and references therein). These results are consistent with findings reported by ^29,30^ for soybean in the US, who reported significant negative effects of temperatures higher than 30°C on yield. Conversely, projected yield increase in the North and East of Europe are mainly associated with positive warming effects in colder regions, where an increase of temperature is expected to have a positive effect on soybean yield because temperatures get closer to the optimum for a number of physiological processes in soybean (Table 2, Table S7). A detailed analysis of the Random Forest algorithm using partial dependence plots relating temperature variables to simulated soybean yield (Figure S7) confirms that model outputs are very consistent with the current knowledge on soybean physiology (Table S7). Indeed, several temperature thresholds established from field experiments are captured in the partial dependence plots: the minimum temperature of 4°C for germination (Figure S7-A), the minimum temperature of 10°C and optimum temperature of 30°C for pollen germination (Figure S7-B-D-E), the maximum temperature of 40°C for crop development pre- and post-anthesis (Figure S7-B-C). Together these results suggest that the soybean yield projections presented in this paper are in line with the current understanding of soybean physiology.

**Figure 3.**
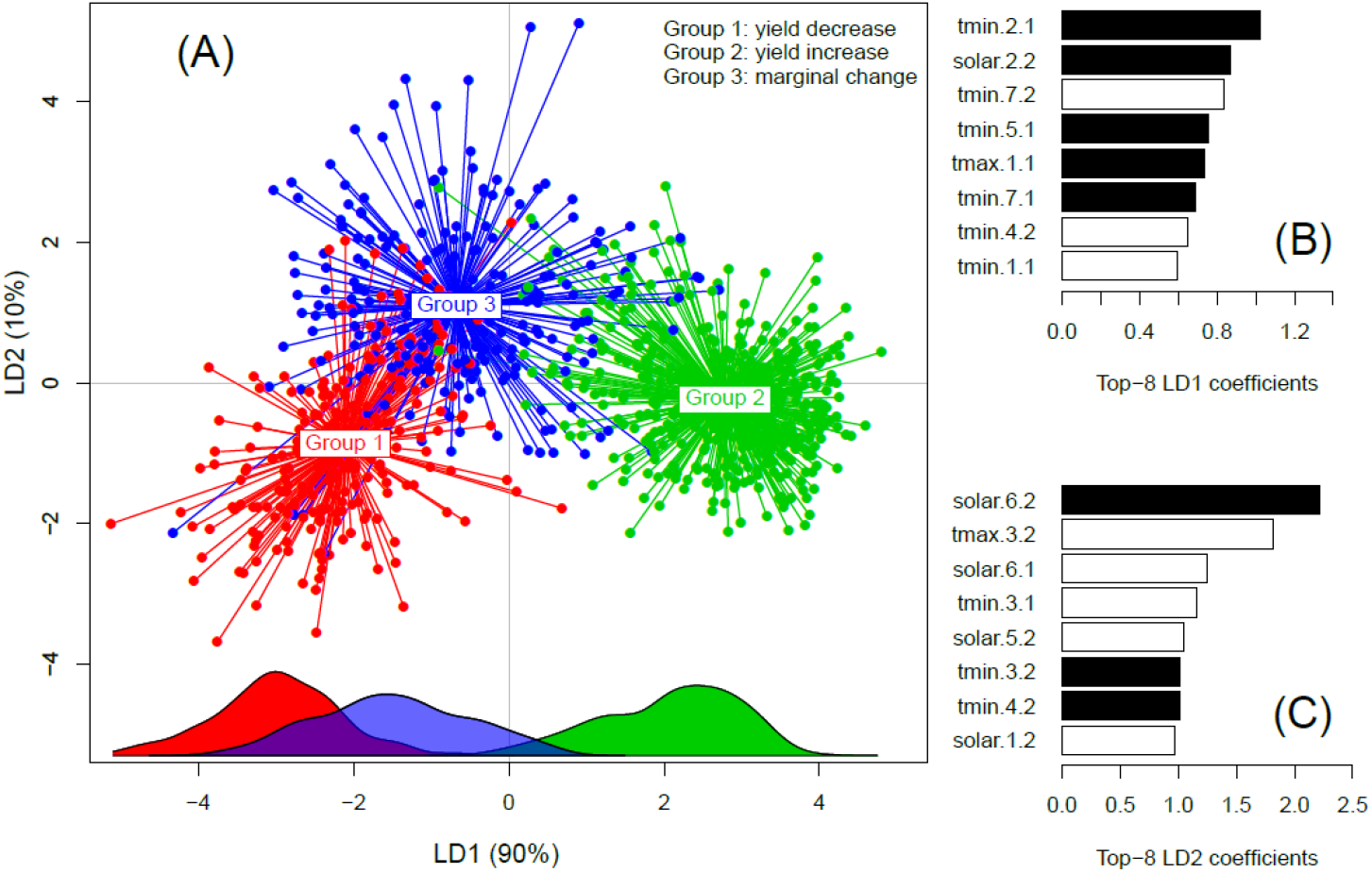
Analysis of climate drivers of projected yield changes by 2050s under RCP 4.5 relative to historical climate. **Panel (A)** shows the linear discriminant analysis (LDA) performed on climate variables for three groups of grid-cells defined by predicted soybean yield change between RCP 4.5 by 2050s and historical climate. Groups of grid-cells are defined as Group 1: yield decrease (projected yield change < - 0.3 t ha^-1^), Group 2: yield increase (projected yield change > 0.3 t ha^-1^), and Group 3: marginal change (yield change between −0.3 and +0.3 t ha^-1^). The plot axes are the two main LDA discriminant functions. Density along the x-axis is shown for each group (same color code applies). See Figure S5 for a map showing the projected yield changes in Europe between RCP 4.5 by 2050s and historical climate. **Panels (B) and (C)** show climate variables contributions to linear discriminant 1 and 2, respectively (the higher the value, the higher the contribution of the corresponding climate input). White bars indicate a positive contribution, and black bars indicate a negative contribution. To improve clarity, only the eight climate variables contributing most to each linear discriminant are shown (see Figure S6 for the contributions of all climate variables). Suffixes to climate variables names indicate, first, the month of the soybean growing season, and second, the time period (“1” standing for historical climate, and “2” standing for the 2050s under RCP 4.5). For example, “tmin.2.1” means “monthly average daily minimum temperature in the second month of the growing season under historical climate”.

**Table 2.** Temperature changes associated with a decrease (Group 1) or a decrease (Group 2) in projected soybean yield under RCP 4.5 by mid-century relative to historical climate. Reported temperature values represent mean values for each group of grid-cells. Group 1: yield decrease (projected yield change < - 0.3 t ha^-1^), Group 2: yield increase (projected yield change > + 0.3 t ha^-1^). Groups are the same than those in the Linear Discriminant Analysis presented in Figure 3, but for clarity, only Group 1 and Group 2 are presented here (see Table S6 for a description of all groups and other climate variable means by group). The GRASP dataset^22^ is used for historical climate, and the median over height Global Circulation Models^25^ is shown for RCP 4.5 by mid-century. Yield projections are performed with the Random Forest algorithm presented in Table 1 and Figure S2.

| Climate variable | Month* | Group 1<br>yield decrease |  | Group 2<br>yield increase |  |
| --- | --- | --- | --- | --- | --- |
|  |  | Historical climate | 2050s (RCP 4.5 ) | Historical climate | 2050s (RCP 4.5 ) |
| Tmax (°C) | 1 | 15,3 | 16,5 | 10,6 | 13,1 |
|  | 2 | 21,1 | 22,6 | 17,6 | 19,6 |
|  | 3 | 25,5 | 27,9 | 21,3 | 23,8 |
|  | 4 | 28,6 | 31,3 | 23,4 | 25,9 |
|  | 5 | 28,0 | 30,9 | 21,7 | 24,2 |
|  | 6 | 23,4 | 26,1 | 16,5 | 18,9 |
|  | 7 | 16,8 | 18,7 | 10,1 | 12,1 |
| Tmin (°C) | 1 | 4,9 | 5,8 | 1,7 | 3,2 |
|  | 2 | 9,7 | 11,1 | 6,9 | 8,8 |
|  | 3 | 13,3 | 15,8 | 10,5 | 13,2 |
|  | 4 | 15,9 | 18,3 | 13,0 | 15,4 |
|  | 5 | 15,3 | 18,0 | 11,7 | 14,0 |
|  | 6 | 11,5 | 13,8 | 7,9 | 9,7 |
|  | 7 | 7,0 | 8,3 | 3,3 | 4,9 |

\* month of the soybean growing season (April to October in Europe)

### Area requirements for 50% and 100% soybean self-sufficiency in Europe

Our projections of soybean suitable area suggest untapped opportunities to increase soybean production in Europe. We estimated the soybean production area required to reach a self-sufficiency level of 50% and 100% based on yield projections presented in Figure 1. A three-step procedure was followed. First, we assumed that soybean could only be grown on current cropland^31^. Under this assumption, soybean cannot be grown in place of permanent pastures, in line with the Common Agricultural Policy of the European Union aiming at their protection^32^. Second, we considered four scenarios for the increase of soybean frequency in crop sequences. In these scenarios, soybean is grown in one year in three, four, five, or six years, which correspond to 33%, 25%, 20%, and 16% cropland area in a grid-cell under soybean, respectively. These scenarios are consistent with observed and recommended soybean frequencies in crop sequences in Europe. Indeed, a 1-in-3 year or 1-in-4 year soybean cultivation is often recommended to limit the risk of disease occurrence^33^ (especially those caused by two fungal pathogens *Sclerotinia sclerotiorum* and *Rhizoctonia solani*^34,35^), although higher frequencies are observed in Europe^33,36^ and other countries^37,38^. Third we assumed that soybean is grown preferably in high-yielding grid-cells. Based on this assumption, soybean areas were allocated to grid-cells ranked in decreasing order of projected yield values until the cumulated production (calculated as the product of area and yield) reached 50% and 100% of current annual soybean consumption of Europe (58 Mt in average over 2009-2013^1^). Results suggest that a self-sufficiency level of 50% (100%) would be achievable in Europe under historical and future climate whatever the frequency of soybean in crop sequences, if 4 to 5% (9 to 12%) of the current European cropland is dedicated to soybean production (Figure 4, Figure S8). For the self-sufficiency level of 50%, this share corresponds to 11.2 to 14.5 Mha or about 2 to 3 times larger than the current European soybean area (5 Mha in 2016^1^). For the self-sufficiency level of 100%, the corresponding values are 25 to 34.4 Mha or about 5 to 7 times larger than the current area (Figure 5A).

**Figure 4.**
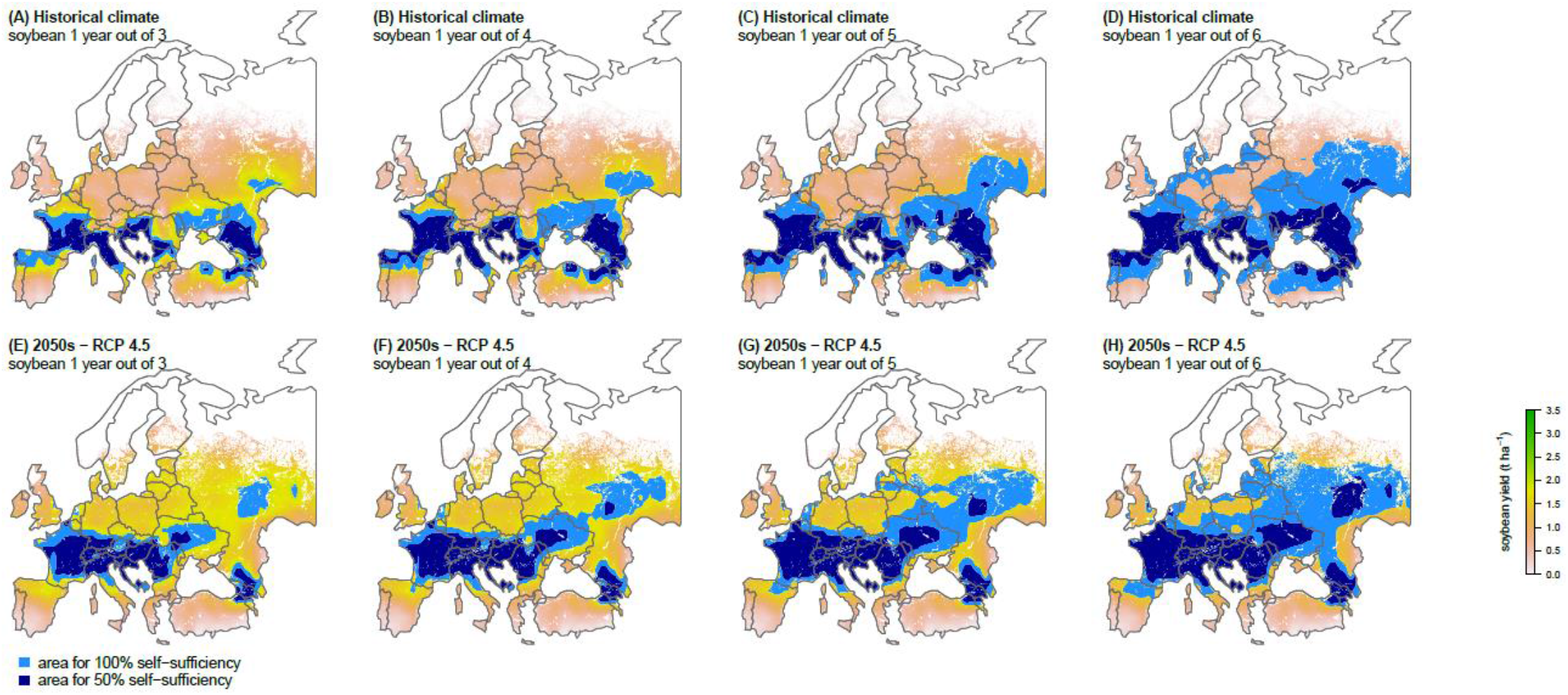
Area requirements for 50% and 100% soybean self-sufficiency in Europe under historical climate (A-D) and by 2050s under RCP 4.5 (E-F). Based on soybean yield projections presented in Figure 1 and assuming various levels of soybean frequency in crop sequences (one year out for three, four, five and six years), soybean areas were allocated to grid-cells ranked in decreasing order of projected yield values until the cumulated production (calculated as the product of area and yield) reached 50% (light blue) and 100% (dark blue) of the current annual soybean consumption of Europe (58 Mt, average 2009-2013). We assume that soybean can only be grown on current cropland^31^, which excludes permanent pastures in line with the Common Agricultural Policy of the European Union aiming at their protection^32^. Background colors indicate projected soybean yield in t ha^-1^ as in Figure 1.

**Figure 5.**
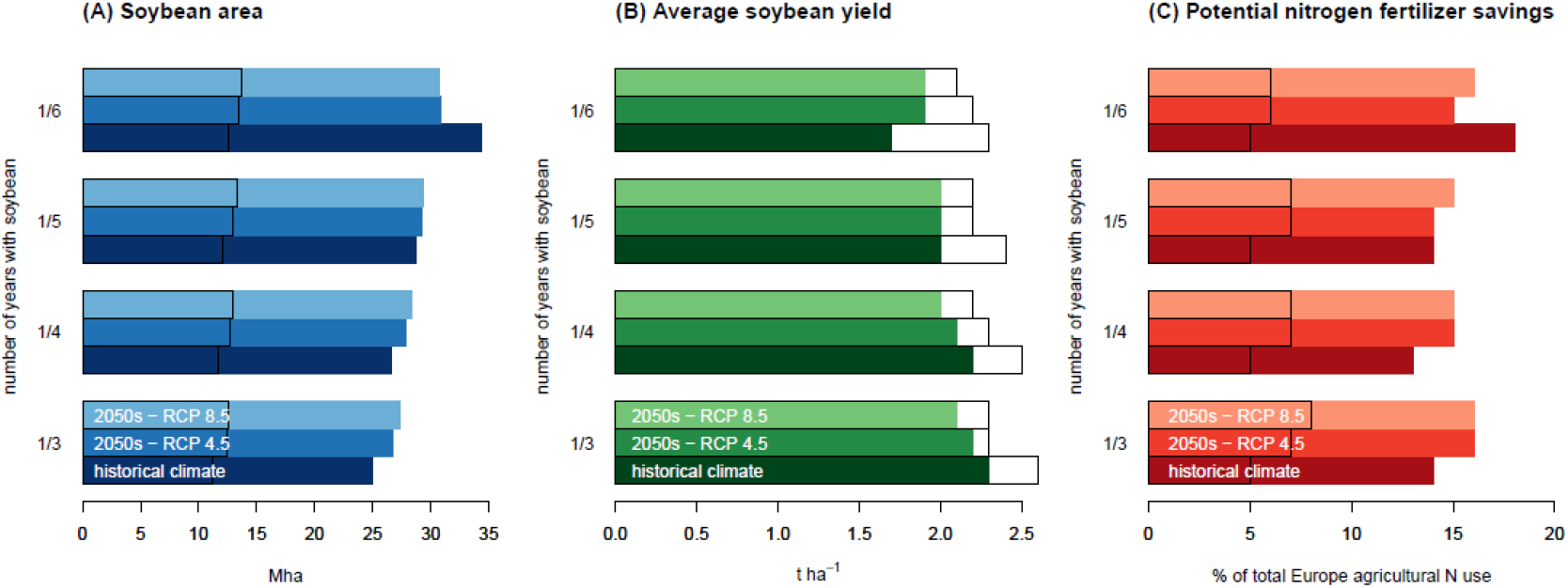
(A) Soybean production area required to reach 50% and 100% soybean self-sufficiency, and associated (B) average soybean yield, and (C) potential N-fertilizer savings, under historical climate (1981-2010) and by 2050s under RCP 4.5 and 8.5. Color-filled bars indicate values corresponding to 100% soybean self-sufficiency, and empty bars with a black border indicate values corresponding to 50% soybean self-sufficiency. Four levels of soybean frequency in crop sequences are evaluated in which soybean is grown in one year in three, four, five, or six years, which correspond to 33%, 25%, 20%, and 16% cropland area in a grid-cell under soybean, respectively. Potential nitrogen fertilizer savings are calculated based on the assumption that soybean is not fertilized with nitrogen and that it will replace N-fertilized crops.

### Potential nitrogen fertilizer savings from soybean expansion

An expansion of soybean area may have some environmental benefits, in particular by reducing nitrogen (N) fertilizer applications. Soybean is an N_2_-fixing crop which is usually fertilized at very low rates or even not fertilized at all with nitrogen, saving N-fertilizer use compared to other crops. Assuming soybean is not fertilized with nitrogen in Europe, and using published global maps of crop-specific N-fertilizer rate^39,40^ (Figures S9 to S11), we estimated N-fertilizer savings from the replacement of fertilized crops (e.g. wheat) by unfertilized soybean. Results show that the extra soybean area needed to reach 50% (100%) self-sufficiency would reduce total N-fertilizer use in Europe by 5 to 8% (13 to 18%) (Figure 5C). This estimate is likely to be conservative because additional N-fertilizer savings could be expected from reducing N-fertilizer rates applied to non-legume crops following soybean in the crop sequence by about 20 kgN ha^-1^, as commonly recommended by agronomists for cereals in relation with the high N content of legume residuals^4,5^. However, an accurate estimation of this positive effect of soybean as a previous crop for cereals would require assumptions about which crops would preferentially be replaced by soybean, which is out of scope of this study but deserves further research.

## Discussion

### Large opportunities to increase soybean production in Europe

Our study shows that soybean suitable area estimated from agro-climatic conditions is much larger than current soybean harvested area in Europe, even under projected climate change. It also suggests that current and future climate would allow Europe to grow enough soybean to reach a self-sufficiency level of 50%, i.e. five time greater than the current level of 10%. Moreover, our projections suggest that achieving 100% self-sufficiency is possible, at least on an agro-climatic point of view. Although the 100% self-sufficiency scenario might not be a realistic target for a number of reasons discussed below, this nevertheless highlights the large opportunities to increase soybean production in Europe. These results have concrete implications for Europe. First, soybean production doesn’t appear to be limited by climate only. Socio-economic factors are currently limiting the development of soybean as well, like low market competitiveness compared to other crops or imported soybean, lack of value chains development, public subsidies in favor of cereals, difficulty to account for non-market environmental benefits of soybean, low information dissemination on best management practices, as previously mentioned by other studies ^2,3,41^. Second, our results show that a shift of soybean suitable areas from the south of Europe towards the north-east of Europe is projected under climate change by the middle of this century, according to the moderate and intense climate change scenarios considered (RCPs 4.5 and 8.5). Therefore, the relative profitability of soybean production in the different countries within Europe might change accordingly. By highlighting regions with high projected soybean yield under both current and future climate, our findings could help policymakers and agrobusinesses reorganizing their production area distribution. This could also be of interest for breeders, who recently started making efforts to create soybean varieties specifically adapted to European conditions, especially high latitudes^42^. In this study, moderate and intense levels of warming were considered. These two emission scenarios were selected to depict maximum impacts of climate change (RCP8.5), with moderate impacts were used as the reference (RCP4.5), although low level of warming (RCP2.6) is worth examining to lead to more climate-policy relevant implications.

### On the environmental impacts of soybean expansion in Europe

Based on a simple assumption that non-fertilized soybean would replace N-fertilized crops, we estimate that the expansion of the soybean area needed to reach 50% and 100% self-sufficiency and the resulting decrease of crop areas receiving high amounts of N-fertilizer would reduce total N-fertilizer use in Europe by 5 to 8% and 13 to 18%, respectively. The excess use of N-fertilizers is widely recognized to be associated with negative impacts on soil, air, and water quality, climate change (through greenhouse gases emissions from manufacture and field application), and biodiversity conservation^43^. Therefore, our results suggest that the reduction in N-fertilizer use resulting from a soybean expansion in Europe could have positive environmental effects. However, further research is necessary to better quantify the possible benefits and side effects. Increasing soybean area might also help controlling pests, diseases, and weeds in European agricultural systems through a diversification of cereal-based intensive cropping systems^6^. For example, it is now well established that diversifying crop sequences help controlling weeds, especially when the diversity of sowing periods (e.g. autumn, spring) in the crop sequence increases^44^. An expansion of the soybean area also questions agricultural water management. Indeed, an increase in soybean acreage will have an impact on water demand, and this impact can be negative or positive depending on which crop is replaced by soybean. As soybean is a summer crop, its cultivation in replacement for a winter crop (e.g. wheat) is expected to increase water demand during summer and impact negatively water resources, especially in Southern Europe^15,27,28^. On the other hand, if soybean replaces another irrigated summer crop such as maize, the demand for summer water should decrease because the amount of water applied is generally higher on irrigated corn than on irrigated soybeans^45^.

### Potential impacts of soybean expansion on land use change in Europe and abroad

Projected shifts in soybean suitable area and possible expansion of soybean area in Europe could have important implications in terms of land use, both in Europe and in major soybean producing countries. For example, if an additional 9 Mha of soybean is grown in place of wheat (this corresponds to area requirements for 50% self-sufficiency), the European wheat production area (60 Mha, average 2013-2017^1^) would be reduced by 15% with strong consequences on wheat production. An expansion of soybean area into previously uncultivated land could have negative impacts on the environment through associated GHGs emissions and potential loss of biodiversity. A large increase in European soybean production would also have potential impacts in other countries. As previously mentioned, Europe currently imports about 90% of its domestic soybean consumption. A large share of these imports come from South America, particularly Argentina, where a link has been established between the deforestation of the Gran Chaco dry tropical forest – a biodiversity hotspot^46^ – and global demand for soybeans^47,48^. Given current soybean yield level in Argentina (2.9 t ha^-1^, average 2013-2017^1^), the additional soybean production needed for Europe to achieve 50% self-sufficiency would represent about 8 Mha of soybean area in Argentina, or more than 40% of the national soybean area in Argentina. Overall, these results suggest that an expansion of the European soybean area could help preventing deforestation in biodiversity hotspots. However, we acknowledge that land use dynamics are difficult to anticipate because of their complexity^49^, and international trade needs be considered through economic modeling.

### On the use of machine learning to model soybean yield

In recent years, machine learning techniques have been successfully applied to predict yields of a variety of crops in different world regions^50–53^. Consistently with those previous studies, the results presented here demonstrate the good predictive ability of the Random Forest algorithm when applied to soybean, with a RMSEP of 0.35 t ha^-1^. Two important results reinforce the reliability of the conclusions drawn from our model. First, the model behavior is very consistent with current knowledge on soybean physiology, as shown by partial dependence plots relating temperature variables to yield (Figure S7, Table S7). This is remarkable as no information was *a priori* included in the model on this point. Second, our projections do not involve any extrapolation of the model beyond the range of data used for training (Figure S4, Table S4). Our machine learning algorithm (RF) has also several advantages compared to standard parametric statistical models. RF does not make any prior assumption on the relationship between soybean yield and climate inputs. RF is able to handle nonlinear effects and complex interactions between climate inputs which would have been difficult to include in standard statistical models. Moreover, with RF, the yield response to climate is data-driven and does not rely on pre-specified equations. The good performance of our model suggests that the combined use of large global climate and yield data sets with machine learning techniques is a promising approach to studying the impact of climate on agricultural production. One key point underlying this is probably the wide range of climate conditions captured in training data. The assessment of transferability in space revealed that our model predictive ability decreased markedly when the distance between training and test grid-cells was higher than 1000 km (Table 1). The global soybean yield dataset we used in this study^22^ contains only a few grid-cells in Europe, which are located in the north of Italy (Figure S12). This low amount of grid-cells located in Europe is consistent with the limited (although expanding) extent of soybean production area in Europe, which was especially true around the year 2000 that corresponds to the time period depicted by the dataset we used^20^. These few grid-cells located in Europe are of great value as they allow capturing local features of soybean yield – climate relationships in our model. However, increasing the number of observed soybean yield data in Europe appears as a great opportunity to improve yield predictions from climate inputs. To this end, databases containing large amounts of experimental data such as the one recently published by Cernay et al.^54^ could be used in the future to update our projections.

### Potential effects of some factors not included in the model

Our predictive model presents some limitations due to the fact that several factors are neglected, in particular day length, soil type, atmospheric CO2 concentration, and shifting growing season due to climate change. Soybean is known to be a short-day plant that needs day length to stay below a given threshold (which depends on the cultivar) to flower at a maximum rate: if day length exceeds this threshold, flowering is delayed and maturity might not be reached before the end the growing season^55^. However, day length has not been included in the predictors of our model because it is a function of latitude^56^, and thus there is a high risk of confounding effects associated with environmental or socio-economic variables correlated with latitude^57^. We believe that the fact that day length was not taken into account in our model is likely to have a small impact on our results for two reasons. First, day length becomes an issue especially at high latitudes, and there is already some evidence from farmer’s fields and field experiments that soybean can be grown at high latitude (up to 55-57 °N) in Europe (Figure S4-A). Second, our model don’t project high yields at latitudes higher than 55-57 °N. Further research will benefit from exploring soybean cultivation at high latitudes with process-based crop models that include the effect of day length on soybean physiology^58^.

Soil type is known to have impacts on crop growth and yield in multiple ways. However, no reliable historical soil dataset is currently available at the global scale for key soil characteristic relevant to crop growth^59,60^. We therefore decided not to include soil type in our model. Nevertheless, we note that some analyses based on global soil datasets suggest no important limitation to root growth in European soils^61^, thus making our yield projections more likely to be pessimistic than optimistic. Additionally, published maps of European soil types^62^ highlight the existence of specific soil types in the North of Europe (e.g. leptosols in Norway, and podzols in Scandinavia), and the North-East of Europe (e.g. albeluvisols in Russia), but these areas are not identified as high-yielding soybean areas in our projections.

In line with previous work, we do not consider the effect of an increase in atmospheric CO2 concentration on soybean yields because this effect is still very uncertain due to many complex interaction mechanisms, and is still widely discussed in the research community^63–68^. However, if an increase in atmospheric CO2 concentration positively affects soybean yield as suggested by the most up-to-date quantitative synthesis of available experimental and modeling studies (which reports an estimated global average yield increase of soybean of +11% for an increase in atmospheric CO2 concentration of +100ppm)^69^, it further supports our findings that large opportunities exist to improve Europe soybean self-sufficiency under climate change. And although it is likely that an increase in atmospheric CO2 concentration will impact absolute yields, there is no evidence that it will change the relative yields of the different geographical regions, and thus the ranking of the grid-cells considered here.

Our projections assume a fixed growing season from April and to October. A shift in the growing season (sowing date and cultivars with different maturity or length of growing cycle) might offer opportunities for crop adaptation to climate change^70^. But investigation of these effects is not straightforward with the RF algorithm we used for projections, and this raises a number of methodological questions that are out of the scope of this paper. For example, a possible caveat of statistical models including RF is that the extent to which a shortened crop duration and associated yield losses under increased temperature conditions is accounted for is unclear. The fixed growing season assumed in this study may be a reason for relatively optimistic soybean yield projections. However, in Northern and Eastern Europe, the positive impact of climate change on yields can be interpreted as the result of a decrease in cold stress, which would compensate for the negative impact of reduced crop growth duration. Here again, further research will benefit from the use of process-based crop models able to capture these processes, to make comparisons with the results presented in this paper.

## Methods

### Soybean yield, irrigated fraction, and climate data

Soybean yields used in this paper are from the global dataset of historical yields updated version ^20,21^. This includes grid-wise soybean yields worldwide with the grid size of 1.125 degree, which covers the period 1981-2010. Yield values reported in this dataset result from the combination of national-scale yield statistics from the FAO, global crop calendars and harvested areas, and satellite-derived net primary production values. Therefore, the grid-cell yields are estimated values resulting from the combination of several sources of information. In this dataset, the soybean harvested areas are those of the year c.a. 2000, and are kept constant (Figure S12). The geographical coverage of soybean harvested area found in^20^ is relatively limited compared to other datasets^40^ because in some parts of the world the crop calendar used to generate grid-cell yield estimates^71^ is missing. Regarding historical climate data, the global retrospective meteorological forcing dataset tailored for agricultural application (GRASP) was used^22^. This dataset contains monthly average of five climatic variables relevant in explaining crop growth and yield: daily maximum and minimum air temperatures at 2m, daily precipitation, daily solar radiation, and daily vapor pressure. These variables are available for the period 1961–2010 at the same spatial resolution as yield data, i.e. a grid size of 1.125 degree. Other meteorological forcing datasets are available^72^, but uncertainties associated with different datasets are small at monthly time scale. The SPAM2005 v3.2 dataset (available at http://mapspam.info/) was used to retrieve irrigated soybean fraction in each grid-cell^73^. This dataset provides the irrigated and rainfed harvested area for a set of crops (including soybean) at the global scale for around the year 2005, at a spatial resolution of 5 arc min (~0.08 degree). These data were regridded to the spatial resolution of the yield data (1.125 degree) using the *projectRaster()* function of the *raster* R package with argument *method* set to *“bilinear”*.

### Data preprocessing

We focus on the major soybean producers representing 91% of the global soybean harvested area, namely Argentina, Brazil, Canada, China, India, Italy, and USA (Figure S12). Information regarding yield data and crop calendars (sowing and harvest dates) – needed to define the growing seasons – may be considered more reliable for major soybean producers than for minor players. Moreover, previous studies suggest that soybean actual yield is close to the estimated yield potential in at least some areas within these countries, e.g. in USA ^74^, Argentina ^75^, and Brazil ^76^. This is of importance because climate effects on soybean yield are easier to detect when non-climatic factors (e.g. sub-optimal management) are not limiting yield. We remove all grid-cells with soybean areas lower than 1% of grid-cell area, a threshold below which we consider soybean production to be too marginal to be included in the analysis. To avoid any confusion with technological progress, soybean yield data are detrended in order to remove the increasing trends of soybean yield time series due to improved cultivars and technological progress ^77,78^. For all grid-cells, yield time series are detrended using a cubic smoothing spline *f(t)* and each yield data is then expressed relatively to the expected yield value in 2010 as *Yd(t) = f(2010) + A(t)*, where *f(2010)* is the smoothing spline yield estimate for the year 2010 (the most recent year available in the yield dataset) and *A(t)* is the yield anomaly *A*(*t*) = *yield*(*t*)-*f*(*t*). Histograms of soybean yields before and after detrending are shown in Figure S13. The soybean growing season is defined country-by-country according to the crop calendars provided by the Agricultural Market Information System (available at: http://www.amis-outlook.org/amis-about/calendars/soybeancal/en/). Based on this source of information, the soybean growing season is considered to range from April to October in China, USA, and Italy, from November to May in Argentina and Brazil, from May to November in Canada, and from June to December in India.

### Adding zero yield data

In order to take into account climate conditions preventing soybean cultivation and leading to zero yields, the yield dataset was expanded by adding grid-cells located in climate zones known to be environmentally unsuitable for crop production, like deserts and arctic areas. Six climate zones from the last version of the Köppen-Geiger climate classification (available at http://koeppen-geiger.vu-wien.ac.at/present.htm, ^79,80^) were selected, and 67 grid-cells were selected at random in each selected climate zone, so that added zero yield values represented 20% of the final dataset. The six selected climate zones are described in Table S8, and a map showing locations of added grid-cells is available in Figure S14. The final dataset includes 30,337 yield data values. This procedure allows us to significantly increase the range of environmental conditions captured in our dataset, which has been shown to have a strong impact on the performances of such models^81^.

### Modeling soybean yield

Detrended soybean yield data is related to 35 climate variables defined at a monthly time step over the seven months of the soybean growing season, plus the fraction of irrigated area, i.e. a total of 36 variables. The 35 climate variables are monthly mean daily minimum and maximum temperatures (*Tmin* and *Tmax*, degree Celsius), monthly total precipitation (*rain*, mm month^-1^), monthly mean daily total solar radiation (*solar*, MJ m^-2^ day^-1^), monthly mean air vapor pressure (VP, hPa). Four different approaches are used to predict yield from the 36 input variables: Artificial Neural Network (ANN), Random Forests (RF), Generalized Additive Model (GAM), and Multiple Linear Regression Model (MLR). All these algorithms were fitted using the *R* software v3.4.0. For ANN we used the *neuralnet()* function of the *neuralnet* package^82^, with one 10-neurons hidden layer and default values for other parameters. RF was fitted with the *ranger()* function of the *ranger* package^83^, with a number of trees set to 500 and default values for other parameters. MLR was fitted with the *glm()* function of *R*, and GAM was fitted with the *gam()* function of the *gam* package^84^.

### Assessing model transferability in time and space

The model predictive ability is first assessed using a bootstrap approach with 25 out-of-bag samples generated by bootstrap, using the *train()* function of the *caret* R package. However, recent articles have highlighted the importance of rigorous cross-validation strategies to ensure that the predictive capacity of a given algorithm is evaluated on data as independent as possible from the data used to train that algorithm^23,24^. Here, we run two cross-validation strategies to assess transferability of the above algorithms in time and space. Transferability in time was assessed by splitting the dataset into two periods in order to assess the ability of each algorithm to predict a period of time different from the one used for the training: 1981-1995, and 1996-2010. In a first step, each algorithm was fitted on 1981-1995 to predict 1996-2010. In a second step, each algorithm was fitted on 1996-2010 to predict 1981-1995. Transferability in space was assessed using a five-step procedure implemented for each algorithm in turn: (i) select a grid-cell at random in the yield database (excluding the zero yield cells), (ii) define 7 buffer zones of different sizes (radius) around the selected grid-cell (radius values are 100 km, 500 km, 1000 km, 1500 km, 2000 km, 2500 km, and 3000 km), (iii) for each buffer zone, remove grid-cells within the buffer zone and fit the algorithm on the rest of the dataset (including added zero yield grid-cells) – note that to avoid any confusion with the size of the training dataset, the training dataset was composed of 700 grid-cells selected at random outside the buffer zone, (iv) predict the 30 years of yield for the grid-cell selected at step (i) considering each buffer zone in turn, (v) compute average error of prediction over years for the selected grid-cell considering each buffer zone in turn. This procedure is repeated over 10 grid-cells selected at random in each country in order to estimate transferability in space for various degree of spatial proximity between the training and test datasets. Assessing transferability in space is key here because of the low number of grid-cells located in Europe in the historical soybean yield dataset (Figure S12). Therefore, making projections of soybean yield in Europe will necessarily imply some degree of transferability in space of the algorithm. In both cases (transferability in space and time), predictive ability was measured by computing the root mean square error of prediction (RMSEP, t ha^-1^), and Nash–Sutcliffe model efficiency (MEF, unitless)^85^. An efficiency of 1 corresponds to a perfect match of modeled to observed data, an efficiency of 0 indicates that predictions are as accurate as the mean of observed data, whereas an efficiency lower than zero occurs when the observed mean is a better predictor than the tested algorithm. The algorithm showing best transferability in time and space (i.e. the lowest RMSEP and highest efficiency) among ANN, RF, GAM, and MLR is used for projections of soybean yield in Europe under current and future climate.

### Projections of soybean suitability area in Europe under current and future climate conditions

We used 16 climate change scenarios consisting of bias-corrected data of eight Global Circulation Models (GCM; GFDL-ESM2M, HadGEM2-ES, IPSL-CM5A-LR, MIROC5, MIROC-ESM, MIROC-ESM-CHEM, MRI-CGCM3, and NorESM1-M, used in the Coupled Model Intercomparison phase 5 (CMIP5)^25^ and two Representative Concentration Pathways (RCPs; 4.5 and 8.5 W m^-2^)^26^. Details on the bias-correction method used here is available in ^86^. Although daily data are available in the bias-corrected GCM outputs, we computed and used monthly data in our analysis. We consider three time periods for projections: 1981-2010 (historical), 2050-2059, and 2090-2099. We present the median predicted soybean yield over the eight GCMs. Soybean growing season used for prediction is April to October. All projections assumed irrigated fraction equals to zero. Projections are shown only on agricultural area (cropland plus pasture), in the year 2000^31^ (Figure S15).

### Analysis of climate drivers of projected yield changes

A Linear Discriminant Analysis (LDA) was performed to identify combinations of climate variables that best discriminate between three groups of grid-cells. These groups of grid-cells are defined as Group 1: yield decrease (projected yield change < - 0.3 t ha^-1^), Group 2: yield increase (projected yield change > +0.3 t ha^-1^), and Group 3: marginal change (yield change between −0.3 and +0.3 t ha^-1^) (see Figure S5 for a map of the geographical repartition of these three groups in Europe). The 0.3 t ha^-1^ threshold was chosen to be higher than the observed interannual variability of soybean yield in Europe, which is 0.2 t ha^-1^ (standard deviation) over the 2000-2014 time period^1^. The LDA was performed with the function *lda()* of the *MASS* R package, with default settings.

### Area requirement for 50% and 100% soybean self-sufficiency in Europe

In average over 2009-2013, Europe domestic supply of soybean was composed of 32 Mt of soybean cake, and 18 Mt of soybean grains (Table S1). Assuming a conversion factor of 0.8 between soybean grains and soybean cake, this is equivalent to a total domestic supply of 58 Mt of soybean grain. We estimate the soybean production area required to reach a self-sufficiency level of 50% and 100% based on yield projections presented in Figure 1. A three-step procedure was followed. First, we assumed that soybean could only be grown on current cropland^31^ (Figure S15 A). Under this assumption, soybean cannot be grown in place of permanent pastures, in line with the Common Agricultural Policy of the European Union aiming at their protection^32^. Second, we considered four scenarios for the increase of soybean frequency in crop sequences. In these scenarios, soybean is grown in one year in three, four, five, or six years, which correspond to 33%, 25%, 20%, and 16% cropland area in a grid-cell under soybean, respectively. Third we assumed that soybean is grown preferably in high-yielding grid-cells. Based on this assumption, soybean areas were allocated to grid-cells ranked in decreasing order of projected yield values until the cumulated production (calculated as the product of area and yield) reached 50% and 100% of current annual soybean consumption of Europe.

### Estimation of potential N-fertilizer savings from soybean expansion

Soybean is an N2-fixing crop which is usually not fertilized with nitrogen, thus generating N-fertilizer savings when it replaces an N-fertilized crop. To estimate potential N-fertilizer savings if soybean production area expanded enough to reach self-sufficiency, we used published global maps of crop-specific N-fertilizer rate and harvested area for wheat, barley, maize, potato, rapeseed, sugarbeet, and sunflower (Figure S9 and S10) ^39,40^. These maps report data for around the year 2000, but represent the most detailed spatially-explicit crop-specific dataset on N-fertilization to date. Crop specific N-fertilizer rate were area-weighted to generate a unique map representing average N-fertilizer rate of major arable crops in Europe (Figure S11). Then, this map was combined with maps of soybean production area required to reach self-sufficiency under the different climate scenarios (Figure 4, and Figure S8) to calculate potential N-savings of soybean area expansion. Finally, calculated potential N-fertilizer savings were compared to total agricultural N use in Europe that is 13.7 Mt N (average in 2009-2013^1^).

## Acknowledgements

This work was supported by the CLAND convergence institute (16-CONV-0003) funded by the French National Research Agency (ANR), by the ACCAF INRA meta-program (COMPROMISE project, COMPROMISE_MP-P10177), and by the LegValue project funded by the European Union’s Horizon 2020 research and innovation programme under grant agreement N°727672. T.I. was partly supported by the Environment Research and Technology Development Fund (S-14) of the Environmental Restoration and Conservation Agency of Japan and Grant-in-Aid for Scientific Research (16KT0036, 17K07984 and 18H02317) of JSPS.

## Author contributions

N.G and D.M. designed research and performed the analysis. T.I. supplied yield and weather data. N.G. wrote the manuscript, with substantial contributions from all co-authors. D.M. initiated research.

## Data availability

The data that support the findings of this study are available from the authors upon request.

## Code availability

R scripts used in this study is available from the authors upon request.

## Footnotes

* « Europe » refers to the FAO category which includes Russia

## References

1. Food and Agriculture Organization of the United Nations. FAOSTAT Statistics Database. (2019). Available at: http://www.fao.org/faostat/en/#data.

2. Magrini, M. B. et al. Why are grain-legumes rarely present in cropping systems despite their environmental and nutritional benefits? Analyzing lock-in in the French agrifood system. Ecol. Econ. 126, 152–162 (2016).

3. Zander, P. et al. Grain legume decline and potential recovery in European agriculture: a review. Agron. Sustain. Dev. 36, (2016).

4. Cernay, C., Makowski, D. & Pelzer, E. Preceding cultivation of grain legumes increases cereal yields under low nitrogen input conditions. Environ. Chem. Lett. 16, 631–636 (2018).

5. Nemecek, T. et al. Environmental impacts of introducing grain legumes into European crop rotations. Eur. J. Agron. 28, 380–393 (2008).

6. Gaba, S. et al. Multiple cropping systems as drivers for providing multiple ecosystem services: from concepts to design. Agron. Sustain. Dev. (2014). doi:10.1007/s13593-014-0272-z

7. Jensen, E. S. et al. Legumes for mitigation of climate change and the provision of feedstock for biofuels and biorefineries. A review. Agronomy for Sustainable Development 32, (2012).

8. Foyer, C. H. et al. Neglecting legumes has compromised human health and sustainable food production. Nat. Plants 2, 16112 (2016).

9. Messina, M., Rogero, M. M., Fisberg, M. & Waitzberg, D. Health impact of childhood and adolescent soy consumption. Nutr. Rev. 75, 500–515 (2017).

10. Zaheer, K. & Akhtar, M. H. An Updated Review of Dietary Isoflavones: Nutrition, Processing, Bioavailability and Impacts on Human Health. Crit. Rev. Food Sci. Nutr. 57, 1280–1293 (2017).

11. Jayachandran, M. & Xu, B. An insight into the health bene fits of fermented soy products. Food Chem. 271, 362–371 (2019).

12. Dold, C. et al. Long-term carbon uptake of agro-ecosystems in the Midwest. Agric. For. Meteorol. 232, 128–140 (2017).

13. Gilmanov, T. G. et al. Productivity and Carbon Dioxide Exchange of Leguminous Crops: Estimates from Flux Tower Measurements. Agron. Journa 106, 545–559 (2014).

14. Urruty, N., Deveaud, T., Guyomard, H. & Boiffin, J. Impacts of agricultural land use changes on pesticide use in French agriculture. Eur. J. Agron. 80, 113–123 (2016).

15. Rüdelsheim, P. L. J. & Smets, G. Baseline information on agricultural practices in the EU Soybean (Glycine max (L.) Merr.). (2012).

16. Martin, N. Domestic soybean to compensate the European protein deficit: illusion or real market opportunity? Oilseeds Fats Crop. Lipids 22, (2015).

17. Krön, M. & Bittner, U. Danube Soya – Improving European GM-free soya supply for food and feed. Oilseeds Fats Crop. Lipids 22, (2015).

18. Venus, T. J., Drabik, D. & Wesseler, J. The role of a German multi-stakeholder standard for livestock products derived from non-GMO feed. Food Policy 78, 58–67 (2018).

19. OECD/FAO. OECD-FAO Agricultural Outlook 2019-2028. (2019).

20. Iizumi, T. et al. Historical changes in global yields: Major cereal and legume crops from 1982 to 2006. Glob. Ecol. Biogeogr. 23, 346–357 (2014).

21. Iizumi, T. et al. Uncertainties of potentials and recent changes in global yields of major crops resulting from census- and satellite-based yield datasets at multiple resolutions. PLoS One 13, e0203809 (2018).

22. Iizumi, T., Okada, M. & Yokozawza, M. A meteorological forcing data set for global crop modeling: Development, evaluation, and intercomparison. J. Geophys. Res. Atmos. Res. 119, 363–384 (2014).

23. Roberts, D. R. et al. Cross-validation strategies for data with temporal, spatial, hierarchical, or phylogenetic structure. Ecography (Cop.). 40, 913–929 (2017).

24. Fourcade, Y., Besnard, A. G. & Secondi, J. Paintings predict the distribution of species, or the challenge of selecting environmental predictors and evaluation statistics. Glob. Ecol. Biogeogr. 27, 245–256 (2018).

25. Taylor, K. e., Stouffer, R. J. & Meehl, G. A. An Overview of CMIP5 and experiment design. Am. Meteorol. Soc. 93, 485–498 (2012).

26. Van Vuuren, D. P. et al. The representative concentration pathways: an overview. Clim. Change 109, 5–31 (2011).

27. Koutroulis, A. G. et al. Freshwater vulnerability under high end climate change. A pan-European assessment. Sci. Total Environ. 614, 271–286 (2018).

28. Iglesias, A. & Garrote, L. Adaptation strategies for agricultural water management under climate change in Europe. Agric. Water Manag. 155, 113–124 (2015).

29. Schlenker, W. & Roberts, M. J. Nonlinear temperature effects indicate severe damages to U.S. crop yields under climate change. Proc. Natl. Acad. Sci. 106, 15594–15598 (2009).

30. Mourtzinis, S. et al. Climate-induced reduction in US-wide soybean yields underpinned by region- and in-season specific responses. Nat. Plants 1, 14026 (2015).

31. Ramankutty, N., Evan, A. T., Monfreda, C. & Foley, J. A. Farming the planet: 1. Geographic distribution of global agricultural lands in the year 2000. Global Biogeochem. Cycles 22, 1–19 (2008).

32. Commision, E. Sustainable land use (greening). Available at: https://ec.europa.eu/info/food-farming-fisheries/key-policies/common-agricultural-policy/income-support/greening_en. (Accessed: 27th February 2020)

33. Đorđević, V., Malidža, G., Vidić, M., Milovac, Ž. & Šeremešić, S. Best practice manual for soya bean cultivation in the Danube region. (Danube Soya, 2016).

34. Hartman, G. L., West, E. D. & Herman, T. K. Crops that feed the World 2. Soybean-worldwide production, use, and constraints caused by pathogens and pests. Food Secur. 3, 5–17 (2011).

35. Pannecoucque, J. et al. Screening for soybean varieties suited to Belgian growing conditions based on maturity, yield components and resistance to Sclerotinia sclerotiorum and Rhizoctonia solani anastomosis group 2-2IIIB. J. Agric. Sci. 1–8 (2018). doi:10.1017/S0021859618000333

36. Grandes cultures biologiques - Les clés de la réussite. (2017).

37. Grassini, P., Specht, J. E., Tollenaar, M., Ciampitti, I. & Cassman, K. G. High-yield maize-soybean cropping systems in the US Corn Belt. in *Crop physiology*. Applications for genetic improvement and agronomy 15–44 (2014).

38. Salembier, C., Elverdin, J. H. & Meynard, J. Tracking on-farm innovations to unearth alternatives to the dominant soybean-based system in the Argentinean Pampa. Agron. Sustain. Dev. 1–10 (2016). doi:10.1007/s13593-015-0343-9

39. Mueller, N. D. et al. Closing yield gaps through nutrient and water management. Nature 490, 254–257 (2012).

40. Monfreda, C., Ramankutty, N. & Foley, J. A. Farming the planet: 2. Geographic distribution of crop areas, yields, physiological types, and net primary production in the year 2000. Global Biogeochem. Cycles 22, 1–19 (2008).

41. Meynard, J. M. et al. Socio-technical lock-in hinders crop diversification in France. Agron. Sustain. Dev. 38, (2018).

42. Kurasch, A. K. et al. Identification of mega-environments in Europe and effect of allelic variation at maturity E loci on adaptation of European soybean. 0000, 1–14 (2017).

43. Houlton, B. Z. et al. A World of Cobenefits: Solving the Global Nitrogen Challenge. Earth’s Futur. 7, 865–872 (2019).

44. Weisberger, D., Nichols, V. & Liebman, M. Does diversifying crop rotations suppress weeds? A meta-analysis. PLoS One 14, 1–12 (2019).

45. Gibson, K. E. B., Gibson, J. P. & Grassini, P. Benchmarking irrigation water use in producer fields in the US central Great Plains. Environ. Res. Lett. 14, 054009 (2019).

46. Nori, J. et al. Protected areas and spatial conservation priorities for endemic vertebrates of the Gran Chaco, one of the most threatened ecoregions of the world. Divers. Distrib. 22, 1212–1219 (2016).

47. Gasparri, N. I., Grau, H. R. & Angonese, J. G. Linkages between soybean and neotropical deforestation: Coupling and transient decoupling dynamics in a multi-decadal analysis. Glob. Environ. Chang. 23, 1605–1614 (2013).

48. Fehlenberg, V. et al. The role of soybean production as an underlying driver of deforestation in the South American Chaco. Glob. Environ. Chang. 45, 24–34 (2017).

49. Meyfroidt, P. et al. Middle-range theories of land system change. Glob. Environ. Chang. 53, 52–67 (2018).

50. Delerce, S. et al. Assessing weather-yield relationships in rice at local scale using data mining approaches. PLoS One 11, (2016).

51. Everingham, Y., Sexton, J., Skocaj, D. & Inman-Bamber, G. Accurate prediction of sugarcane yield using a random forest algorithm. Agron. Sustain. Dev. 36, (2016).

52. Jeong, J. H. et al. Random Forests for Global and Regional Crop Yield Predictions. PLoS One 11, e0156571 (2016).

53. Partridge, T. F. et al. Mid-20th century warming hole boosts US maize yields. Environ. Res. Lett. 14, 114008 (2019).

54. Cernay, C., Pelzer, E. & Makowski, D. A global experimental dataset for assessing grain legume production. Sci. data 3, 160084 (2016).

55. Setiyono, T. D. et al. Understanding and modeling the effect of temperature and daylength on soybean phenology under high-yield conditions. F. Crop. Res. 100, 257–271 (2007).

56. Forsythe, W. C., Rykiel, E. J., Stahl, R. S., Wu, H. i. & Schoolfield, R. M. A model comparison for daylength as a function of latitude and day of year. Ecol. Modell. 80, 87–95 (1995).

57. Hafner, S. Trends in maize, rice, and wheat yields for 188 nations over the past 40 years: A prevalence of linear growth. Agric. Ecosyst. Environ. 97, 275–283 (2003).

58. Schoving, C. et al. Combining Simple Phenotyping and Photothermal Algorithm for the Prediction of Soybean Phenology: Application to a Range of Common Cultivars Grown in Europe. Front. Plant Sci. 10, 1–14 (2020).

59. Folberth, C. et al. Uncertainty in soil data can outweigh climate impact signals in global crop yield simulations. Nat. Commun. 7, 1–13 (2016).

60. Guilpart, N. et al. Rooting for food security in Sub-Saharan Africa. Environ. Res. Lett. 12, 114036 (2017).

61. Shangguan, W., Hengl, T., Mendes de Jesus, J., Yuan, H. & Da, Y. Mapping the global depth to bedrock for land surface modeling. J. Adv. Model. Earth Syst. 9, 65–88 (2017).

62. Jones, A., Montanarella, L. & Jones, R. Soil atlas of Europe. (2005).

63. Cober, E. R. & Morrison, M. J. Soybean Yield and Seed Composition Changes in Response to Increasing Atmospheric CO 2 Concentration in Short-Season Canada. Plants 8, 250 (2019).

64. Thomey, M. L., Slattery, R. A., Bernacchi, C. J., Köhler, I. H. & Ort, D. R. Yield response of field-grown soybean exposed to heat waves under current and elevated [CO2]. Glob. Chang. Biol. 25, 4352–4368 (2019).

65. Vera, U. M. R., Bernacchi, C. J., Siebers, M. H. & Ort, D. R. Canopy warming accelerates development in soybean and maize, offsetting the delay in soybean reproductive development by elevated CO2 concentrations. Plant Cell Environ. 41, 2806–2820 (2018).

66. Li, Y. et al. Elevated CO2 Increases Nitrogen Fixation at the Reproductive Phase Contributing to Various Yield Responses of Soybean Cultivars. Front. Plant Sci. 8, 1546 (2017).

67. Gray, S. B. et al. Intensifying drought eliminates the expected benefits of elevated carbon dioxide for soybean. Nat. Plants 2, 16132 (2016).

68. Xu, G. et al. Soybean grown under elevated CO2 benefits more under low temperature than high temperature stress: Varying response of photosynthetic limitations, leaf metabolites, growth, and seed yield. J. Plant Physiol. 205, 20–32 (2016).

69. Makowski, D., Marajo-Petitzon, E., Durand, J. L. & Ben-Ari, T. Quantitative synthesis of temperature, CO2, rainfall, and adaptation effects on global crop yields. Eur. J. Agron. 115, (2020).

70. Mourtzinis, S., Specht, J. E. & Conley, S. P. Defining Optimal Soybean Sowing Dates across the US. Sci. Rep. 9, 2800 (2019).

71. Sacks, W. J., Deryng, D., Foley, J. A. & Ramankutty, N. Crop planting dates: an analysis of global patterns. Glob. Ecol. Biogeogr. 19, 607–620 (2010).

72. Ruane, A. C., Goldberg, R. & Chryssanthacopoulos, J. Climate forcing datasets for agricultural modeling: Merged products for gap-filling and historical climate series estimation. Agric. For. Meteorol. 200, 233–248 (2015).

73. You, L., Wood, S., Wood-Sichra, U. & Wu, W. Generating global crop distribution maps: From census to grid. Agric. Syst. 127, 53–60 (2014).

74. Grassini, P. et al. Soybean yield gaps and water productivity in the western U.S. Corn Belt. F. Crop. Res. 179, 150–163 (2015).

75. Merlos, F. A. et al. Potential for crop production increase in Argentina through closure of existing yield gaps. F. Crop. Res. 184, 145–154 (2015).

76. Sentelhas, P. C. et al. The soybean yield gap in Brazil - Magnitude, causes and possible solutions for sustainable production. J. Agric. Sci. 153, 1394–1411 (2015).

77. Grassini, P., Eskridge, K. M. & Cassman, K. G. Distinguishing between yield advances and yield plateaus in historical crop production trends. Nat. Commun. 4, 2918 (2013).

78. Ray, D. K., Ramankutty, N., Mueller, N. D., West, P. C. & Foley, J. a. Recent patterns of crop yield growth and stagnation. Nat. Commun. 3, 1293 (2012).

79. Kottek, M., Grieser, C., Beck, C., Rudolf, B. & Rubel, F. World Map of the Köppen-Geiger climate classification updated. Meteorol. Zeitschrift 15, 259–263 (2006).

80. Rubel, F., Brugger, K., Haslinger, K. & Auer, I. The climate of the European Alps: Shift of very high resolution Köppen-Geiger climate zones 1800-2100. Meteorol. Zeitschrift 26, 115–125 (2017).

81. Dupin, M. et al. Effects of the training dataset characteristics on the performance of nine species distribution models: Application to Diabrotica virgifera virgifera. PLoS One 6, (2011).

82. Günther, F. & Fritsch, S. neuralnet: Training of neural networks. R J. 2, 30–38 (2010).

83. Wright, M. N. & Ziegler, A. ranger: A Fast Implementation of Random Forests for High Dimensional Data in C++ and R. J. Stat. SoftwareSoftware 77, 1–17 (2017).

84. Hastie, T. gam: Generalized Additive Models, R Package, version 0.98. R Found. Stat. Comput. Vienna, Austria. (2013).

85. Wallach, D., Makowski, D., Jones, J. W. & Brun, F. Working with dynamic crop models: methods, tools and examples for agriculture and environment. (Academic Press., 2018).

86. Minamikawa, K., Fumoto, T., Iizumi, T., Cha-un, N. & Pimple, U. Prediction of future methane emission from irrigated rice paddies in central Thailand under different water management practices. Sci. Total Environ. 566–567, 641–651 (2016).

