## Supplementary Information for "Data-driven yield projections suggest large opportunities to improve Europe’s soybean self-sufficiency under climate change"

**Content of the supplementary information**

### Supplementary figures for main text

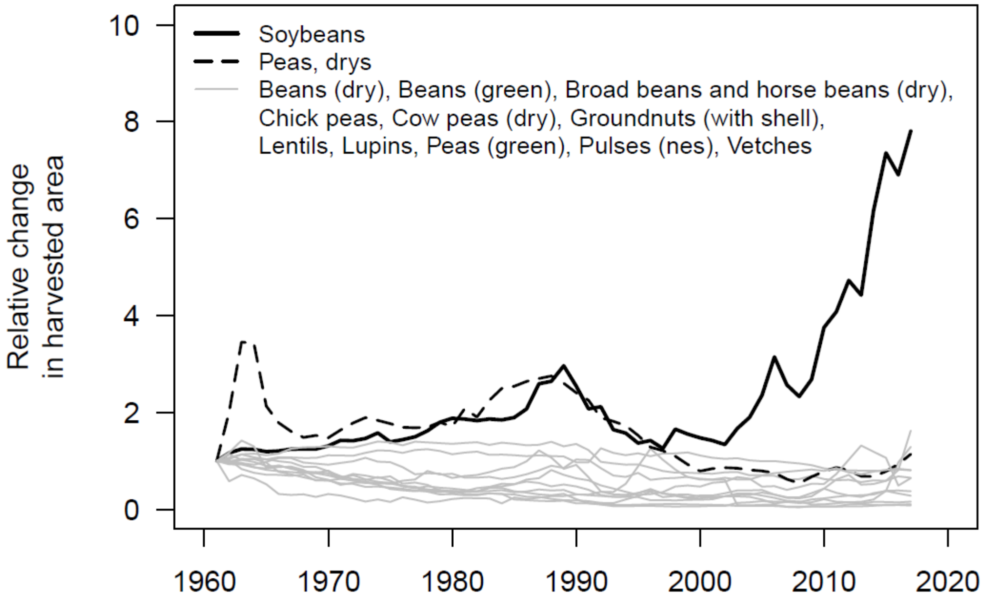

**Figure S1. Historical trends in legumes harvested area in Europe from 1961 to 2017.** Areas are expressed relatively to 1961 (set equal to 1). *Source: FAOSTAT*[1].

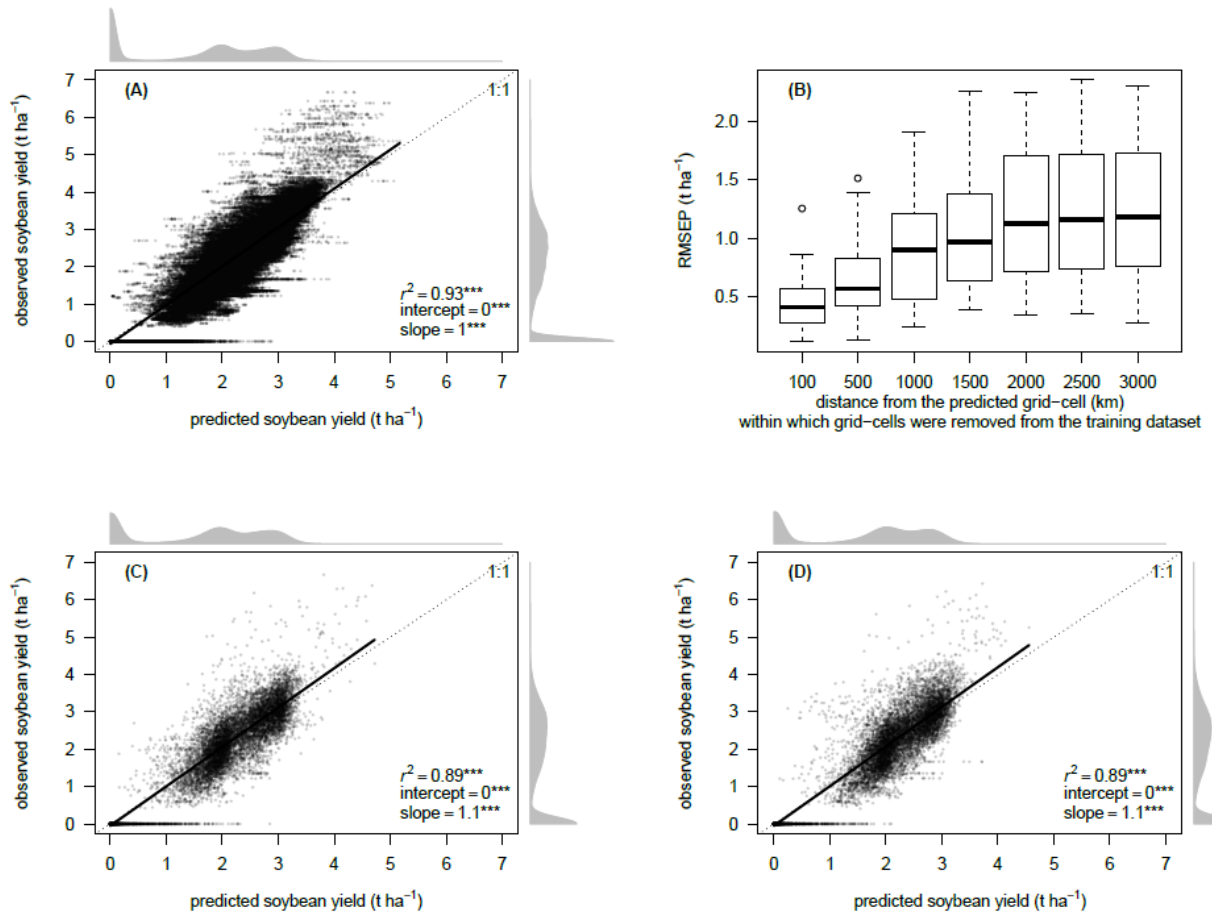

**Figure S2. Assessment of the Random Forest algorithm.** (A) The model is first evaluated using a classical bootstrap approach with 25 resamplings. (B) Model transferability in space is then evaluated by ensuring a minimum spatial distance between training and test datasets. Finally, model transferability in time is assessed in (C) where model is fitted on 1981-1995 to predict 1996-2010, and in (D) where model is fitted on 1996-2010 to predict 1981-1995. RMSEP: root mean square error of prediction. Boxplot in panel (B) shows median (center line), 1$^{st}$ and 3$^{rd}$ quartiles (box limits), and 1.5 times the interquartile range (whiskers). Linear regression outputs are shown on panels (A), (C), and (D), as well as marginal distributions of observed and predicted soybean yields (in grey). Dotted lines represent the 1:1 line. In order to extend the range of climate conditions captured by the model and to capture climate conditions leading to zero yield, additional data points were randomly sampled in climate zones known to be unsuitable for soybean production (e.g. deserts and arctic areas) and added to the dataset with their yield value set to zero.

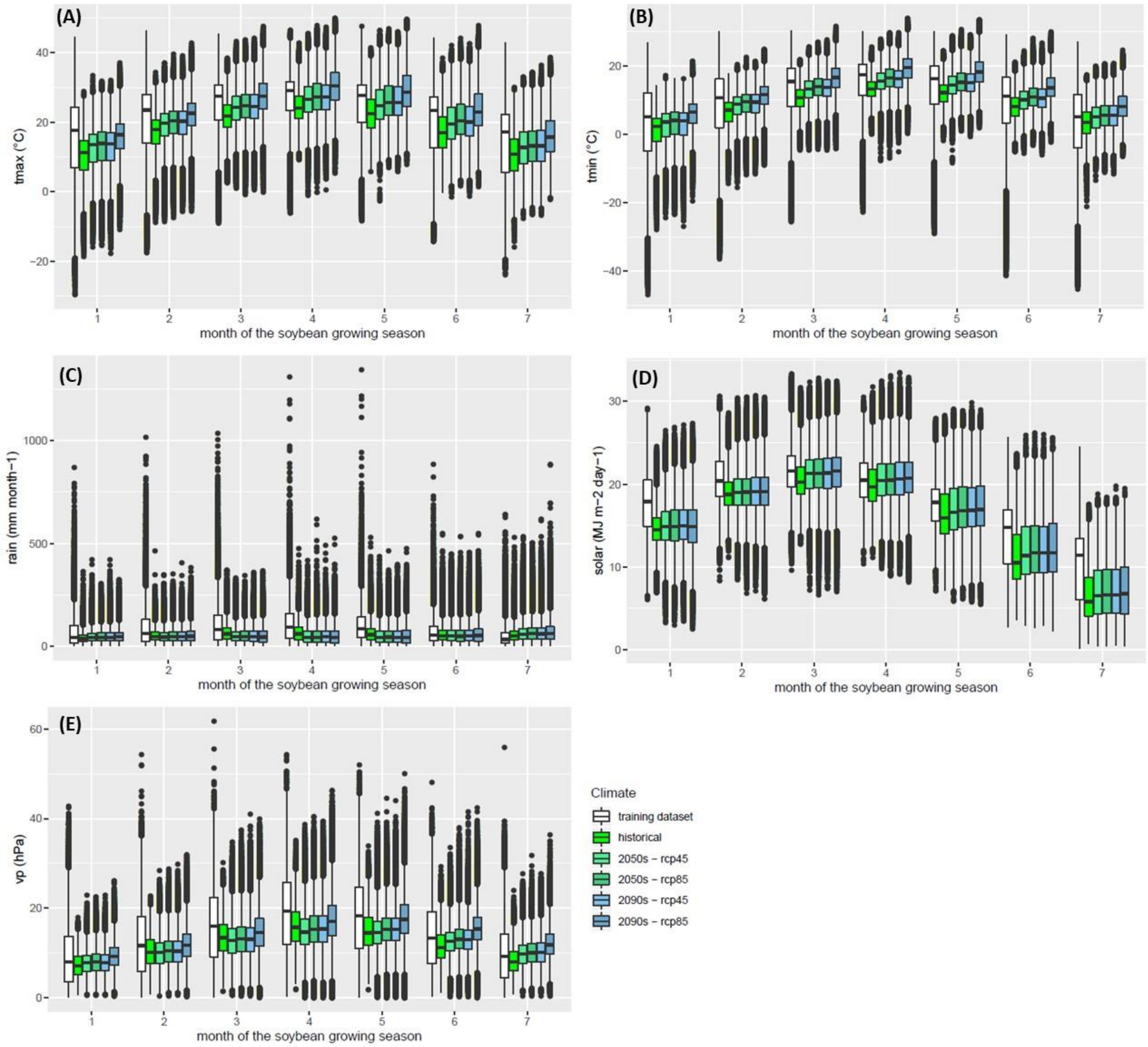

**Figure S3. Comparison of observed ranges of climate variables in the dataset used for training the Random Forest algorithm and the datasets used to make soybean yield projections in Europe under historical and future climate scenarios.** (A) Monthly average of daily maximum temperature. (B) Monthly average of daily minimum temperature. (C) Rainfall. (D) Solar radiation. (E) Vapour pressure. The GRASP dataset[2] is used for model training at the global scale ("training dataset" in the color key) and to make projections of soybean yield in Europe under historical climate (1981-2010) ("historical" in the color key). Then eight Global Circulation Models[3] (see methods) are considered for future climate scenarios, for two periods of time (2050s and 2090s) and two RCPs (RCP 4.5 and RCP 8.5).

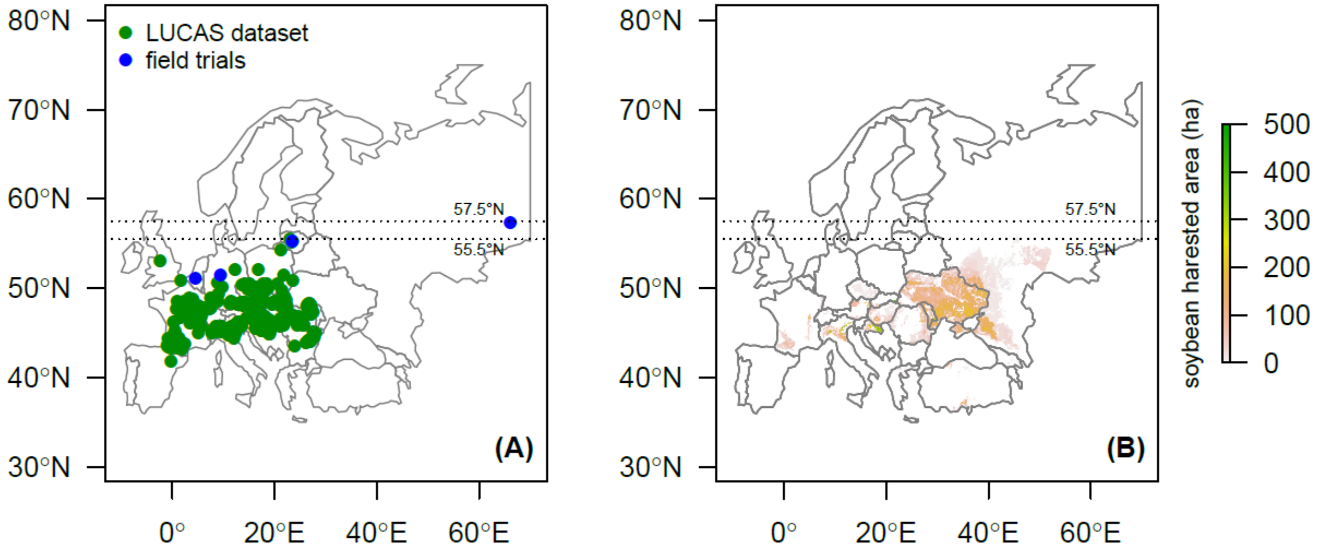

**Figure S4. Evidence map of soybean cultivation at high latitude in Europe.** (A) Locations of farmers soybean fields in 2018 reported by the Land Use and Coverage Area frame Survey (LUCAS) of the European Union (green dots), and locations of some selected published field experiments reporting satisfactory soybean yield (from 2.5 to 4 t $ha^{-1}$) at high latitudes in Europe (blue dots). See Table S4 for details about those field experiments. (B) Harvested soybean area in Europe according to the SPAM2010 model. Sources: the LUCAS dataset is available at https://esdac.jrc.ec.europa.eu/projects/lucas ; and the SPAM2010 dataset is available at: https://www.mapspam.info/ .

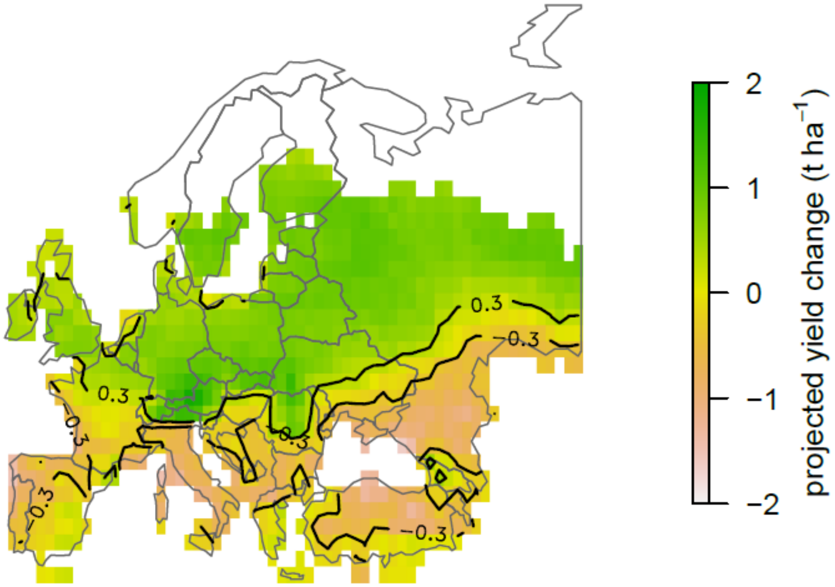

**Figure S5. Projected soybean yield change between RCP 4.5 by mid-century and historical climate.** Projections are performed with the Random Forest algorithm and the GRASP dataset[2] for historical climate (1981-2010), and 8 Global Circulation Models for future climate scenarios[3]. Grid-cells below -0.3 t $ha^{-1}$ correspond to Group 1 (yield decrease) in the Linear Discriminant Analysis (see Figure 3 and Table 2), while grid-cells higher than +0.3 t $ha^{-1}$ correspond to Group 2 (yield increase), and grid-cells between -0.3 and +0.3 t $ha^{-1}$ correspond to Group 3 (marginal change).

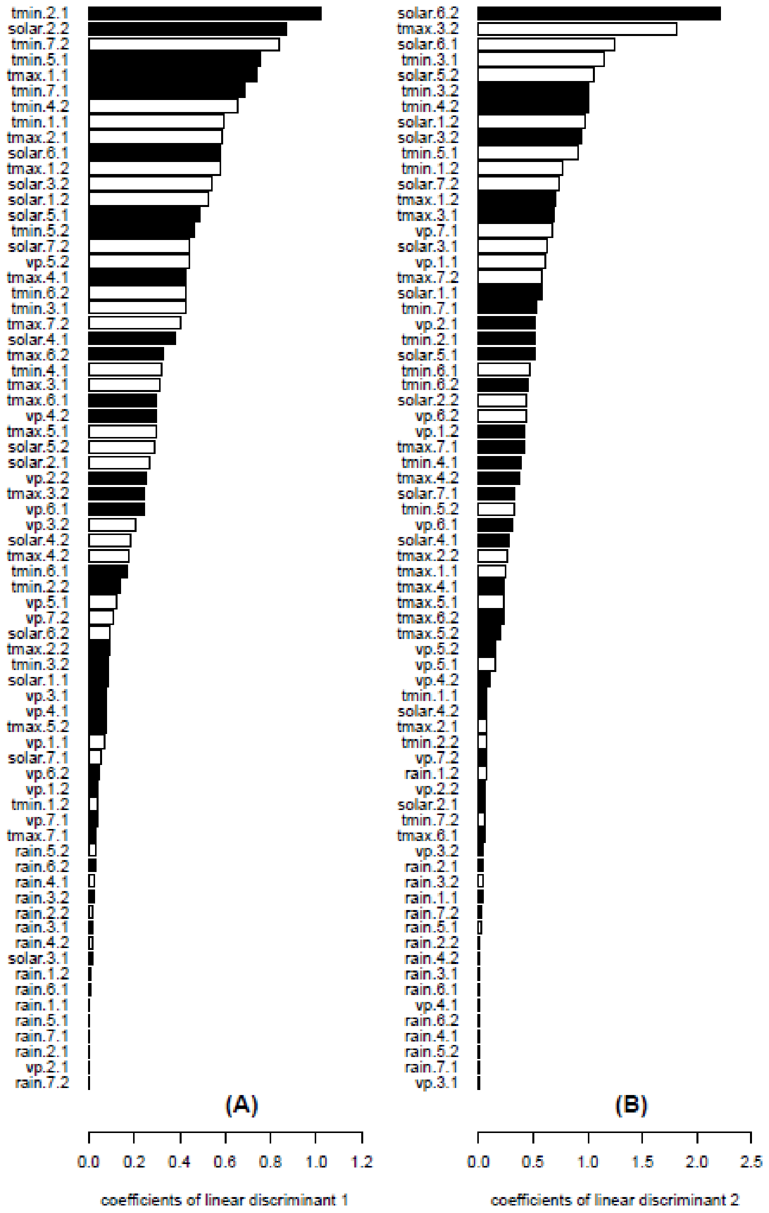

**Figure S6. Climate variables contributions to linear discriminant 1 (panel A) and 2 (panel B).** A linear discriminant analysis (LDA) was performed on climate variables for three groups of grid-cells defined by predicted soybean yield change between RCP 4.5 by 2050s and historical climate. Groups of grid-cells are defined as Group 1: yield decrease (projected yield change < - 0.3 t $ha^{-1}$), Group 2: yield increase (projected yield change > 0.3 t $ha^{-1}$), and Group 3: marginal change (yield change between -0.3 and +0.3 t $ha^{-1}$). White bars indicate a positive contribution, and black bars indicate a negative contribution. The higher the value, the higher the contribution of the corresponding climate input. Suffixes to climate variables names indicate, first, the month of the soybean growing season, and second, the time period ("1" standing for historical climate, and "2" standing for the 2050s under RCP 4.5). For example, "tmin.2.1" means "monthly average daily minimum temperature in the second month of the growing season under historical climate".

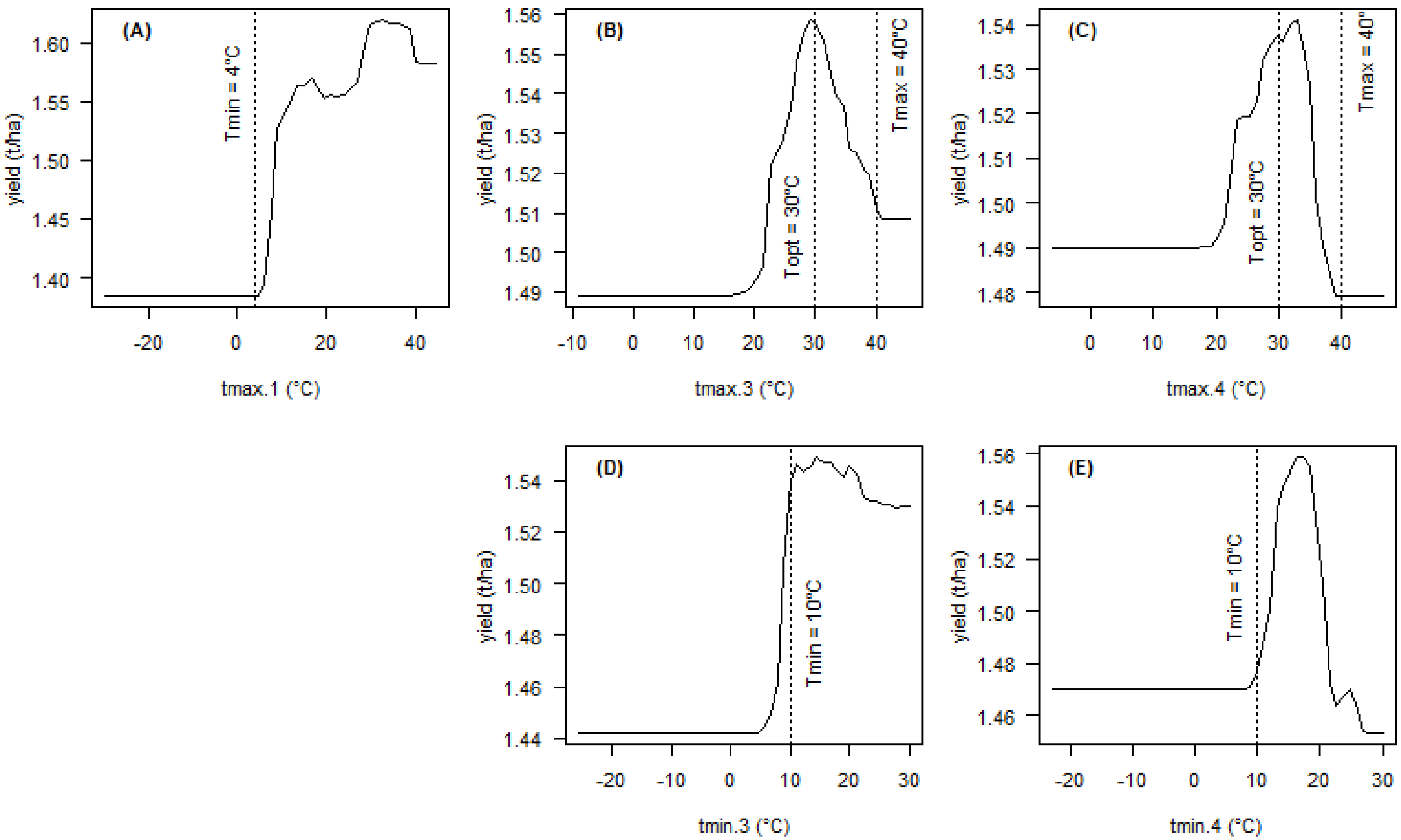

**Figure S7. Partial dependence plots for selected variables of the Random Forest algorithm.** (A) Daily maximum temperature in the first month of the growing season (tmax.1). (B) Daily maximum temperature in the third month of the growing season (tmax.3). (C) Daily maximum temperature in the fourth month of the growing season (tmax.4). (D) Daily minimum temperature in the third month of the growing season (tmin.3). (E) Daily minimum temperature in the third month of the growing season (tmin.3). Dotted lines indicate some cardinal temperatures extracted from the literature to show consistency between the Random Forest algorithm and the current knowledge of soybean physiology. Readers are referred to Table S7 for details about those cardinal temperatures.

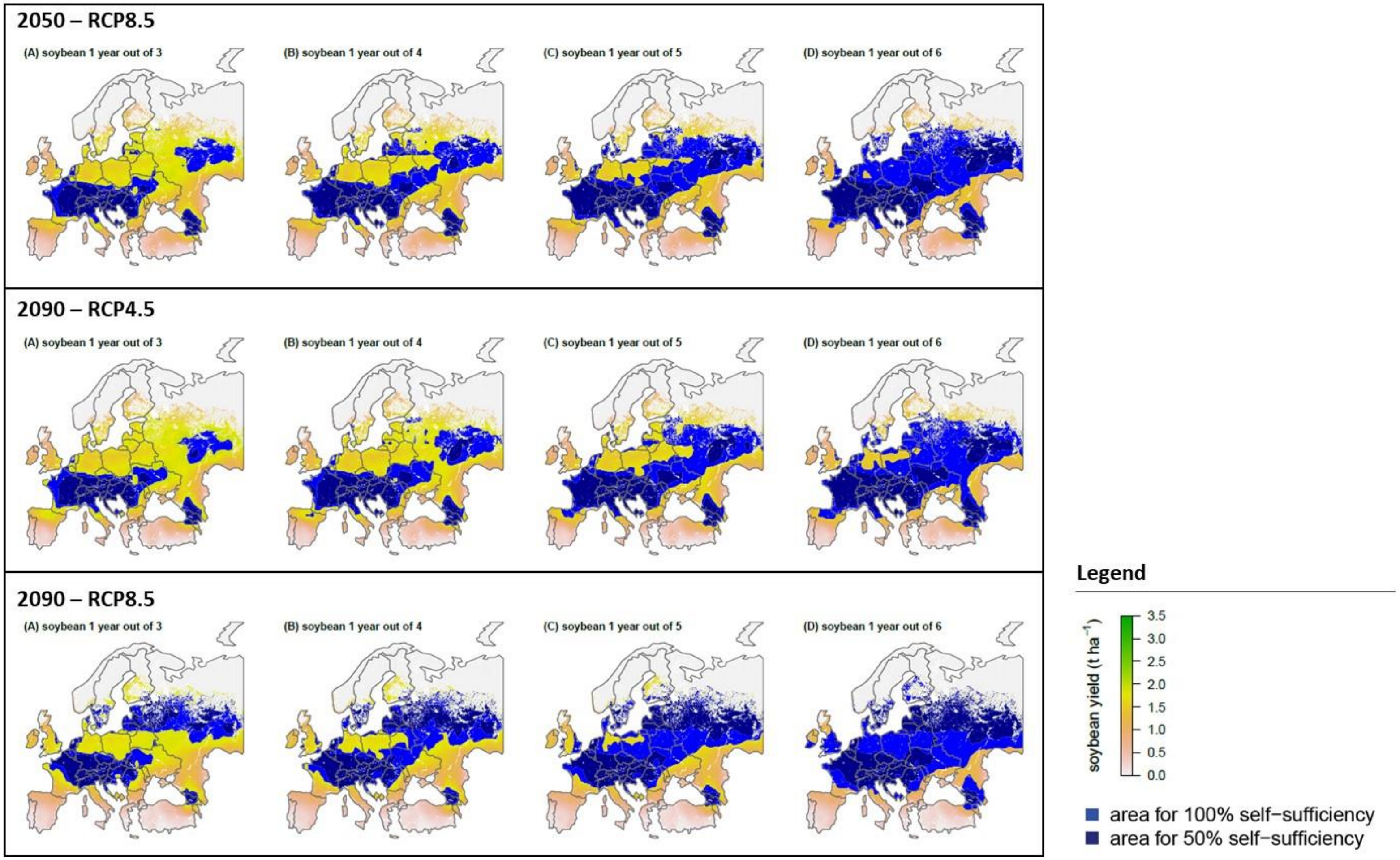

**Figure S8. Area requirements for 50% and 100% soybean self-sufficiency in Europe by 2050s under RCP 4.5 and by 2090s under RCP 4.5 and 8.5.** Based on soybean yield projections presented in Figure 1 and assuming various levels of soybean frequency in crop sequences (one year out for three, four, five and six years), soybean areas were allocated to grid-cells ranked in decreasing order of projected yield values until the cumulated production (calculated as the product of area and yield) reached 50% (light blue) and 100% (dark blue) of the current annual soybean consumption of Europe (58 Mt, average 2009-2013). We assume that soybean can only be grown on current cropland[31], which excludes permanent pastures in line with the Common Agricultural Policy of the European Union aiming at their protection[32]. Background colors indicate projected soybean yield in t $ha^{-1}$ as in Figure 1.

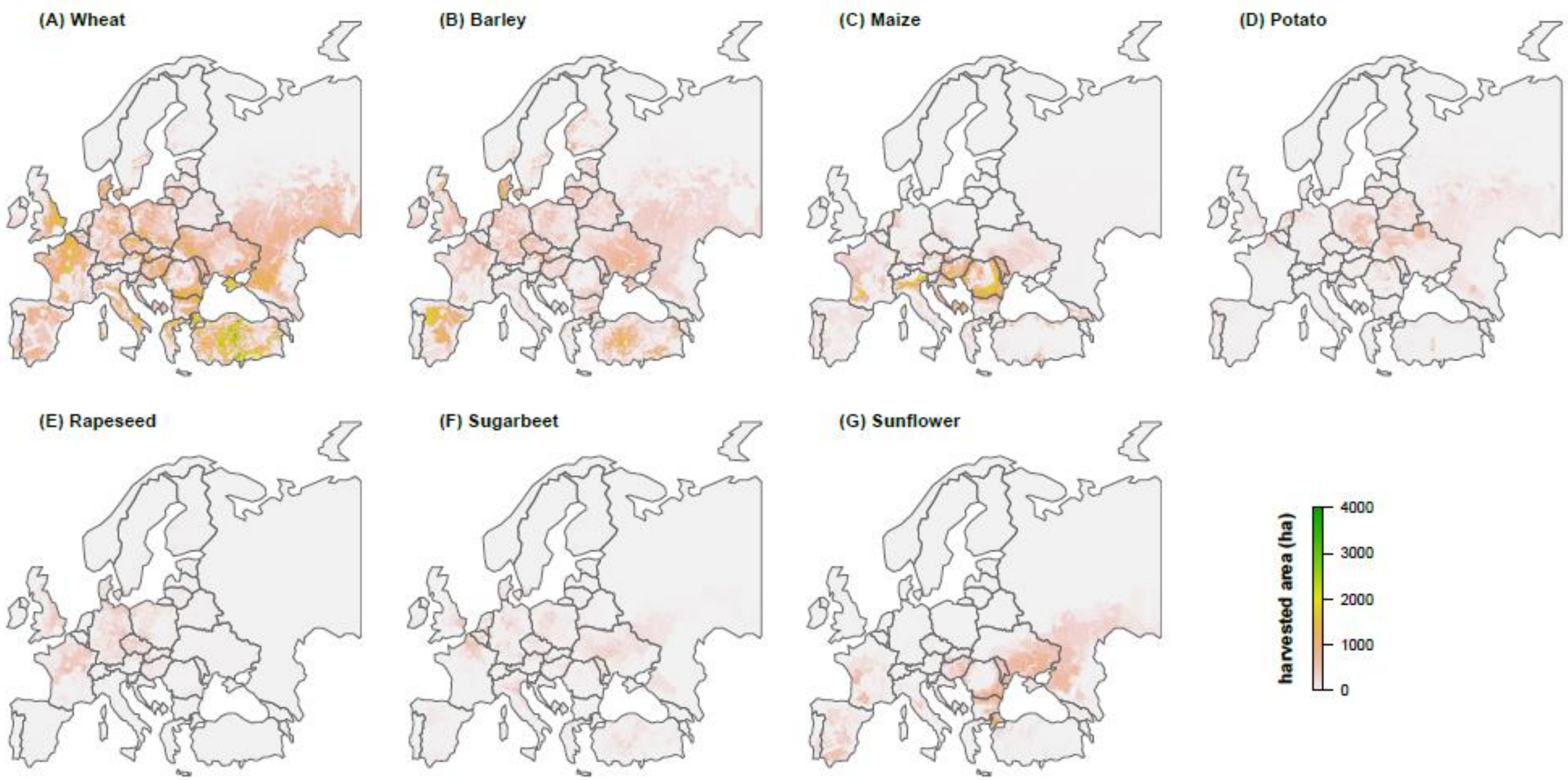

Figure S9. Harvested area maps of (A) wheat, (B) barley, (C) maize, (D) potato, (E) rapeseed, (F) sugarbeet, (G) sunflower in Europe around the year 2000[4].

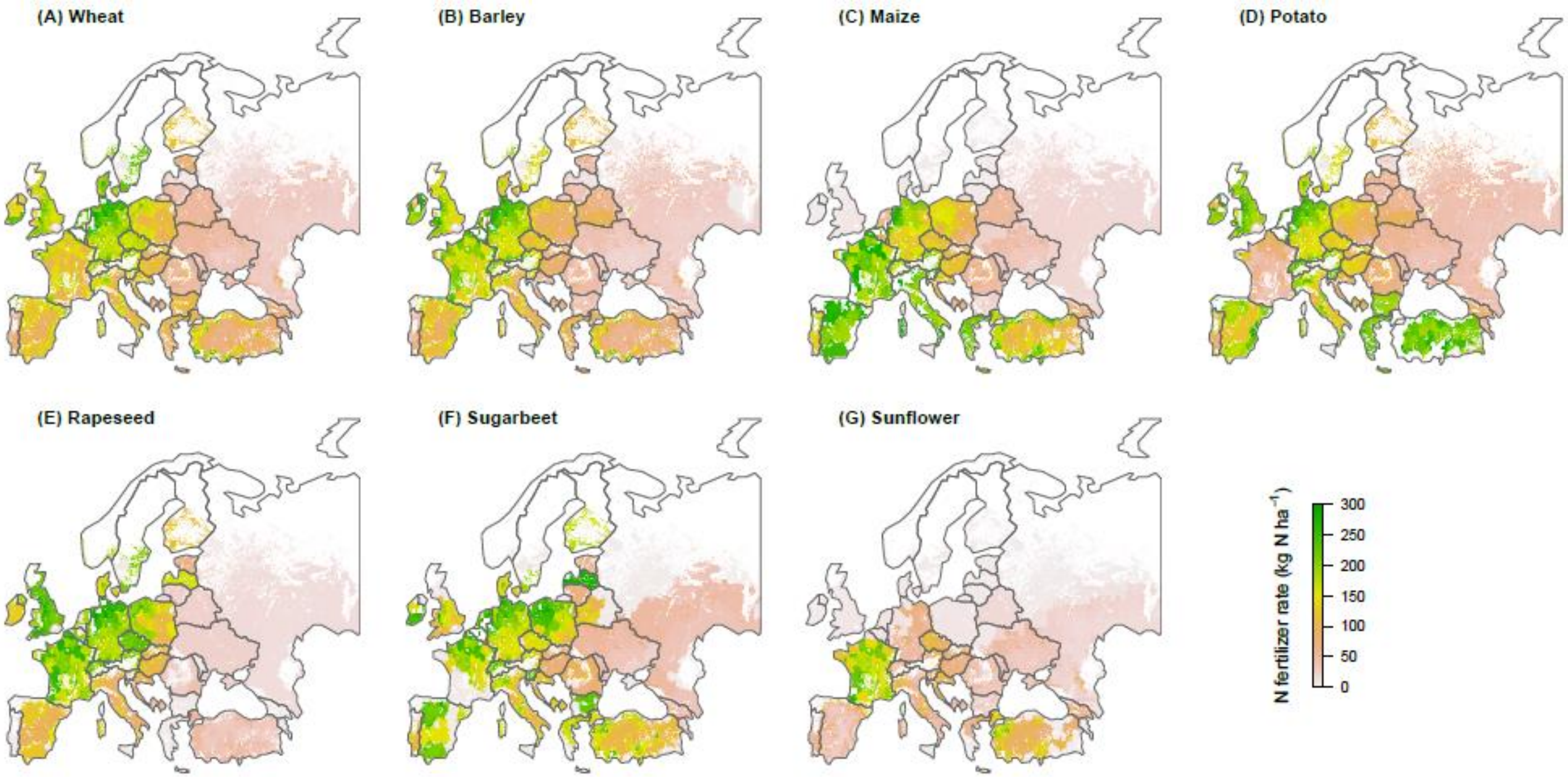

Figure S10. Crop specific nitrogen fertilizer rate for (A) wheat, (B) barley, (C) maize, (D) potato, (E) rapeseed, (F) sugarbeet, (G) sunflower in Europe around the year 2000[5].

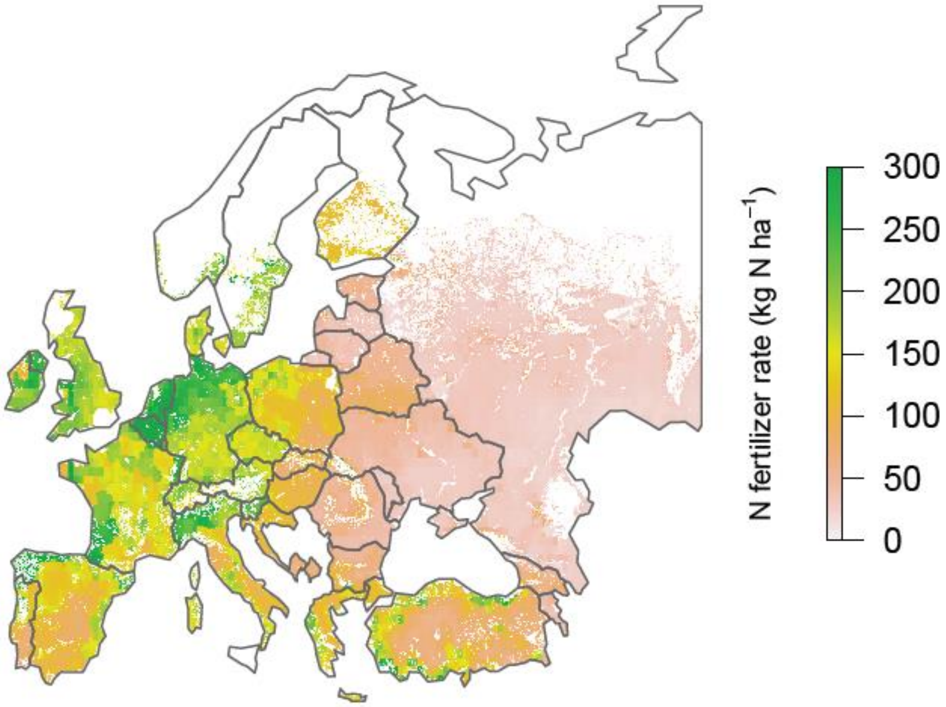

**Figure S11. Average nitrogen fertilizer rate of major crops in Europe around the year 2000.** Map shows the area-weighted nitrogen fertilizer rate based on crop specific nitrogen fertilizer rate and harvested area maps presented in Figure S9 and S10, respectively.

### Supplementary tables for main text

**Table S1. Soybean food balance in Europe (average 2007-2013).** Soybean cake quantity is expressed in soybean grain equivalent assuming that 1 Mt of soybean grain gives 0.8 Mt of soybean cake. *Source: FAOSTAT*[1].

| | Soybean cake | Soybean grain | Total (grain eq.) |
|---|---|---|---|
|  | ---------------------- | *Mt* | ---------------------- |
| **Domestic supply quantity** | 32 | 18 | 58 |
| **Domestic supply breakdown by source** | | | |
| Production | 12 | 5 | 20 |
| Export Quantity | 9 | 3 | 14 |
| Import Quantity | 29 | 16 | 52 |
| Stock Variation | 0 | 0 | 0 |
| **Domestic supply utilization** | | | |
| Feed | 32 | 2 | 42 |
| Food | - | 0 | 0 |
| Processing | - | 16 | 16 |
| Losses | - | 0 | 0 |
| Seed | - | 0 | 0 |
| Other uses | 0 | 0 | 0 |

**Table S2. Changes in soybean suitable area due to climate change in Europe.** All values are in Mha. Percentage change relative to historical climate is indicated in parenthesis.

| | | Area with predicted soybean yield | | |
|---|---|---|---|---|
| **Climate scenario** | | **≥ 1.5 t ha$^{-1}$** | **≥ 2 t ha$^{-1}$** | **≥ 2.5 t ha$^{-1}$** |
| Historical climate (1981-2010) | | 198 | 98 | 31 |
| 2050s | RCP 4.5 | 271 (+37%) | 89 (-10%) | 10 (-69%) |
| | RCP 8.5 | 265 (+34%) | 78 (-20%) | 8 (-75%) |
| 2090s | RCP 4.5 | 287 (+45%) | 96 (-2%) | 12 (-62%) |
| | RCP 8.5 | 256 (+29%) | 45 (-54%) | 0.1 (-100%) |

**Table S3. Percentage of climate data used for soybean yield projections that fall outside of the observed range of climate data used for model training.** Rain: rainfall. Solar: solar radiation. Tmax: daily maximum temperature. Tmin: daily minimum temperature. VP: vapour pressure. For example, 0.13 % of the solar radiation data in month 4 of the soybean growing season under historical climate for the period 1981-2010 fall outside of the observed range for that variable in the dataset used for model training.

| | Month* | Rain | Solar | Tmax | Tmin | VP |
|---|---|---|---|---|---|---|
| **Historical climate** | | | | | | |
| 1981-2010 | 1 | 0 | 0 | 0 | 0 | 0 |
| | 2 | 0 | 0 | 0 | 0 | 0 |
| | 3 | 0 | 0 | 0 | 0 | 0 |
| | 4 | 0 | 0.13 | 0 | 0 | 0 |
| | 5 | 0 | 0.03 | 0 | 0 | 0 |
| | 6 | 0 | 0 | 0 | 0 | 0 |
| | 7 | 0 | 0 | 0 | 0 | 0 |
| **RCP 4.5** | | | | | | |
| 2050s | 1 | 0 | 0.10 | 0 | 0 | 0 |
| | 2 | 0 | 0 | 0 | 0 | 0 |
| | 3 | 0 | 0.01 | 0 | 0 | 0 |
| | 4 | 0 | 0.35 | 0 | 0 | 0.01 |
| | 5 | 0 | 0.13 | 0 | 0 | 0 |
| | 6 | 0 | 0.00 | 0 | 0 | 0 |
| | 7 | 0 | 0 | 0 | 0 | 0 |
| 2090s | 1 | 0 | 0.14 | 0 | 0 | 0 |
| | 2 | 0 | 0.01 | 0 | 0 | 0 |
| | 3 | 0 | 0.02 | 0 | 0 | 0 |
| | 4 | 0 | 0.41 | 0 | 0 | 0.01 |
| | 5 | 0 | 0.18 | 0 | 0 | 0 |
| | 6 | 0 | 0.00 | 0 | 0 | 0 |
| | 7 | 0 | 0 | 0 | 0 | 0 |
| **RCP 8.5** | | | | | | |
| 2050s | 1 | 0 | 0.15 | 0 | 0 | 0 |
| | 2 | 0 | 0.01 | 0 | 0 | 0 |
| | 3 | 0 | 0.04 | 0 | 0 | 0 |
| | 4 | 0 | 0.33 | 0.01 | 0 | 0.01 |
| | 5 | 0 | 0.14 | 0 | 0 | 0.01 |
| | 6 | 0 | 0.00 | 0 | 0 | 0 |
| | 7 | 0 | 0 | 0 | 0 | 0 |
| 2090s | 1 | 0 | 0.34 | 0 | 0 | 0 |
| | 2 | 0 | 0.05 | 0 | 0 | 0 |
| | 3 | 0 | 0.06 | 0.03 | 0.01 | 0 |
| | 4 | 0 | 0.35 | 0.18 | 0.20 | 0.05 |
| | 5 | 0 | 0.13 | 0.06 | 0.27 | 0.02 |
| | 6 | 0 | 0 | 0.02 | 0.01 | 0.01 |
| | 7 | 0 | 0 | 0 | 0 | 0 |

*month of the soybean growing season (April to October in Europe)

Table S4. Details of the selected published soybean field experiments at high latitude in Europe reported in Figure S4.

| **Latitude** | **Country** | **Reported soybean yield** | **References** |
|---|---|---|---|
| 57.35 ° N | Russia | 4 t $ha^{-1}$ | Kühling et al. (2018). Soybeans in high latitudes : effects of Bradyrhizobium inoculation in Northwest Germany and southern West Siberia. Organic Agriculture, 8(2), 159–171.[6] |
| 55.24 ° N | Lithuania | 2.5 - 3 t $ha^{-1}$ | Kadžiulienė et al. (2016). Legumes for sustainability of agroecosystems. In Z. Gaile & Š. Dace (Eds.), 20th Baltic Agronomy Forum (p. 56). Jelgava, Latvia.[7] |
| 51.40 ° N | Germany | 2.5 - 3 t $ha^{-1}$ | Zimmer et al. (2016). Effects of soybean variety and Bradyrhizobium strains on yield, protein content and biological nitrogen fixation under cool growing conditions in Germany. European Journal of Agronomy, 72, 38–46. [8] |
| 51.10 ° N | Belgium | 2.5 - 3 t $ha^{-1}$ | Pannecoucque et al. (2018). Screening for soybean varieties suited to Belgian growing conditions based on maturity, yield components and resistance to Sclerotinia sclerotiorum and Rhizoctonia solani anastomosis group 2-2IIIB. Journal of Agricultural Science, 1–8. [9] |

**Table S5. Confusion matrix of the Linear Discriminant Analysis.** The linear discriminant analysis was performed on climate on climate variables for three groups of grid-cells defined by predicted soybean yield change between RCP 4.5 by 2050s and historical climate. Groups are defined as Group 1: yield decrease (projected yield change < - 0.3 t $ha^{-1}$), Group 2: yield increase (projected yield change > 0.3 t $ha^{-1}$). Group 3: marginal change (yield change between -0.3 and +0.3 t $ha^{-1}$). Overall accuracy (fraction of correct predictions) is 89%.

| | **Predicted group** | | |
|---|---|---|---|
| **Actual Group** | **Group 1** | **Group 2** | **Group 3** |
| **Group 1** | 207 | 1 | 43 |
| **Group 2** | 0 | 491 | 20 |
| **Group 3** | 26 | 25 | 188 |

**Table S6. Analysis of climate variables associated with a decrease (Group 1), increase (Group 2), and marginal change (Group 3) in projected soybean yield under RCP 4.5 by mid-century relative to historical climate.** Reported values of climate variables represent mean values for each group of grid-cells calculated with a Linear Discriminant Analysis (Figure 3). Groups of grid-cells are defined as Group 1: yield decrease (projected yield change < - 0.3 t $ha^{-1}$), Group 2: yield increase (projected yield change > 0.3 t $ha^{-1}$), and Group 3: marginal change (yield change between -0.3 and +0.3 t $ha^{-1}$). The GRASP dataset[2] is used for historical climate, and the median over height Global Circulation Models[3] is shown for RCP 4.5 by mid-century. Yield projections are performed with the Random Forest algorithm presented in Table 1 and Figure S2.

| | | **Group 1 (yield decrease)** | | **Group 2 (yield increase)** | | **Group 3 (marginal change)** | |
|---|---|---|---|---|---|---|---|
| **Climate variable** | **Month*** | Historical climate | 2050s | Historical climate | 2050s | Historical climate | 2050s |
| **Tmax (°C)** | 1 | 15,3 | 16,5 | 10,6 | 13,1 | 15,2 | 16,7 |
| | 2 | 21,1 | 22,6 | 17,6 | 19,6 | 20,9 | 22,4 |
| | 3 | 25,5 | 27,9 | 21,3 | 23,8 | 25,0 | 27,5 |
| | 4 | 28,6 | 31,3 | 23,4 | 25,9 | 27,8 | 30,4 |
| | 5 | 28,0 | 30,9 | 21,7 | 24,2 | 27,1 | 29,9 |
| | 6 | 23,4 | 26,1 | 16,5 | 18,9 | 22,8 | 25,3 |
| | 7 | 16,8 | 18,7 | 10,1 | 12,1 | 16,6 | 18,6 |
| **Tmin (°C)** | 1 | 4,9 | 5,8 | 1,7 | 3,2 | 5,0 | 6,1 |
| | 2 | 9,7 | 11,1 | 6,9 | 8,8 | 9,5 | 10,9 |
| | 3 | 13,3 | 15,8 | 10,5 | 13,2 | 13,1 | 15,5 |
| | 4 | 15,9 | 18,3 | 13,0 | 15,4 | 15,5 | 17,8 |
| | 5 | 15,3 | 18,0 | 11,7 | 14,0 | 15,0 | 17,5 |
| | 6 | 11,5 | 13,8 | 7,9 | 9,7 | 11,5 | 13,6 |
| | 7 | 7,0 | 8,3 | 3,3 | 4,9 | 7,1 | 8,6 |
| **Rain (mm $month^{-1}$)** | 1 | 48,1 | 42,8 | 37,5 | 44,9 | 44,9 | 43,4 |
| | 2 | 49,2 | 41,4 | 53,7 | 48,0 | 44,5 | 41,9 |
| | 3 | 49,4 | 31,8 | 72,2 | 52,2 | 48,9 | 35,4 |
| | 4 | 31,6 | 19,3 | 75,1 | 52,1 | 39,3 | 26,9 |
| | 5 | 32,1 | 15,7 | 68,5 | 52,6 | 39,3 | 25,9 |
| | 6 | 42,6 | 27,6 | 56,9 | 56,9 | 45,7 | 36,4 |
| | 7 | 51,4 | 47,6 | 53,5 | 64,9 | 54,3 | 53,1 |
| **Solar (MJ $m^{-2}$ $day^{-1}$)** | 1 | 16,9 | 17,6 | 14,2 | 14,5 | 16,7 | 17,4 |
| | 2 | 21,0 | 21,4 | 18,4 | 18,6 | 20,7 | 21,1 |
| | 3 | 23,4 | 23,9 | 19,4 | 20,8 | 22,9 | 23,4 |
| | 4 | 23,7 | 24,2 | 18,9 | 19,8 | 22,9 | 23,4 |
| | 5 | 20,8 | 21,4 | 15,5 | 16,1 | 19,9 | 20,5 |
| | 6 | 15,8 | 16,6 | 10,3 | 11,0 | 15,1 | 15,8 |
| | 7 | 10,2 | 11,0 | 5,6 | 6,3 | 9,7 | 10,4 |
| **VP (hPa)** | 1 | 9,2 | 9,1 | 7,4 | 7,9 | 9,3 | 9,3 |
| | 2 | 12,4 | 12,1 | 10,7 | 10,3 | 12,3 | 12,0 |
| | 3 | 15,5 | 14,7 | 14,2 | 13,1 | 15,4 | 14,5 |
| | 4 | 17,8 | 16,5 | 16,3 | 15,1 | 17,8 | 16,6 |
| | 5 | 17,0 | 15,7 | 15,2 | 15,0 | 17,3 | 16,2 |
| | 6 | 13,9 | 14,3 | 11,7 | 12,8 | 14,1 | 14,7 |
| | 7 | 10,3 | 11,8 | 8,4 | 9,9 | 10,8 | 12,2 |

* month of the soybean growing season (April to October in Europe)

**Table S7. Soybean cardinal temperatures.** All values are expressed in degree Celsius (°C).

| Process | Tmin | Topt | Tmax | References |
|---|---|---|---|---|
| Germination | 4 | 30 | 40 | [10] |
| Pollen germination | 10-13 | 28.5 - 30 | 47 | [11,12] |
| Leaf photosynthesis | 5 | 36 | 50 | [13,14] |
| Crop development (phenology) | | | | |
| Pre-anthesis | 5-7.6 | 30-31.5 | 40-45 | [15] |
| Post-anthesis | 3.6-6 | 23-26 | 39-40 | [15–19] |

### Supplementary Figures and Tables for methods

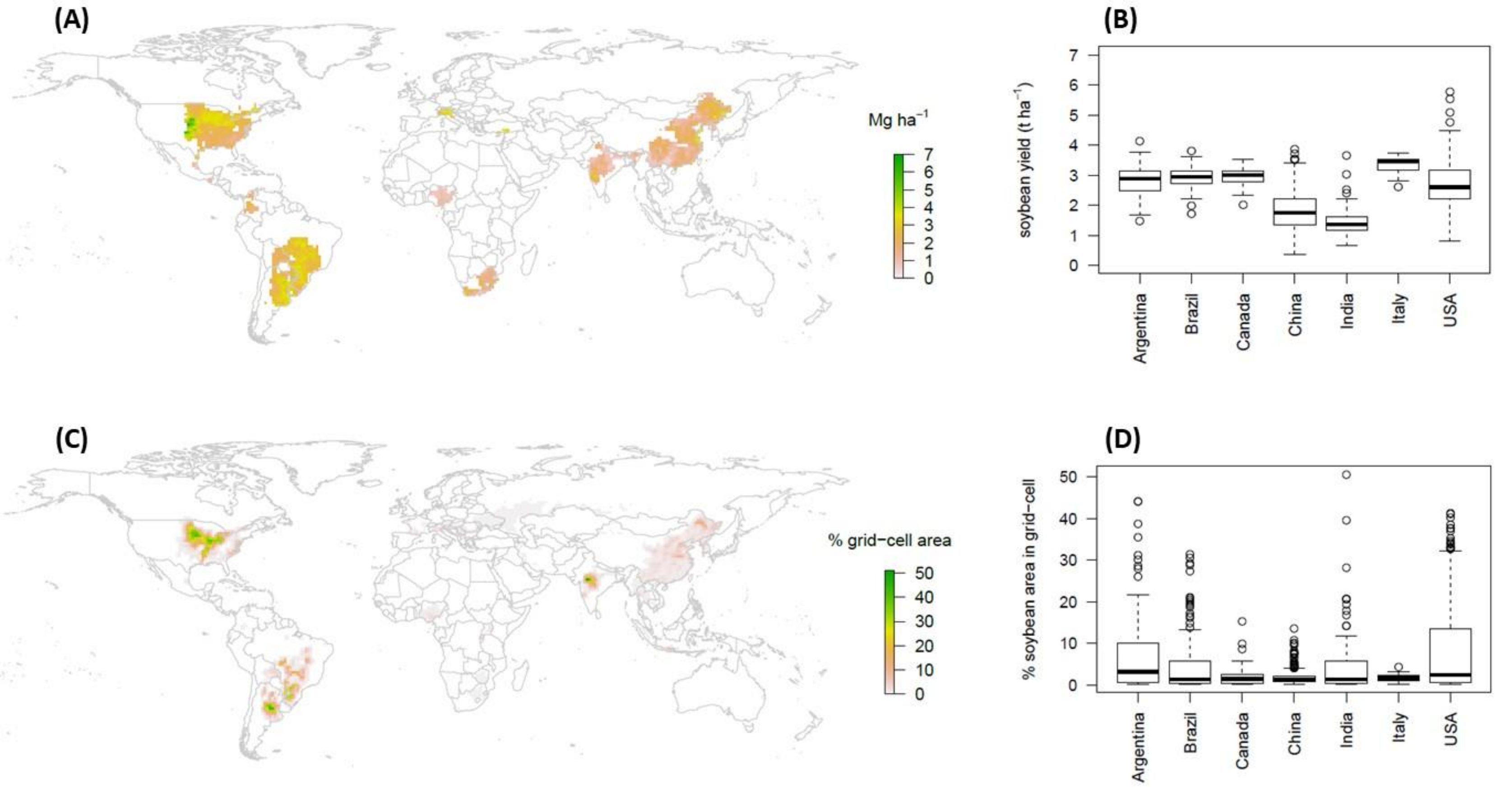

**Figure S12. Soybean yield and harvested area from the global dataset of historical yields**[20]. (A) Global map of soybean yield in 2010. (B) Soybean yield in 2010 in selected countries. (C) Global map of soybean harvested area. (D) Soybean harvested area in selected countries.

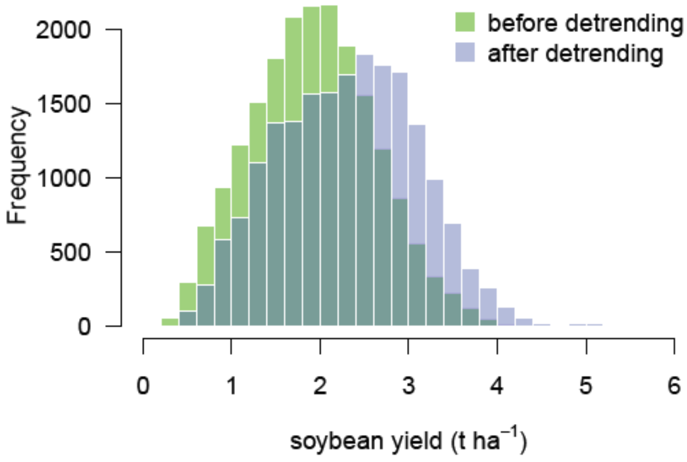

**Figure S13. Histograms of soybean yield before and after detrending.** Detrending is performed in order to remove the increasing trends of soybean yield time series due to improved cultivars and technological progress. Data shown in this Figure are for Argentina, Brazil, Canada, China, India, Italy, and USA.

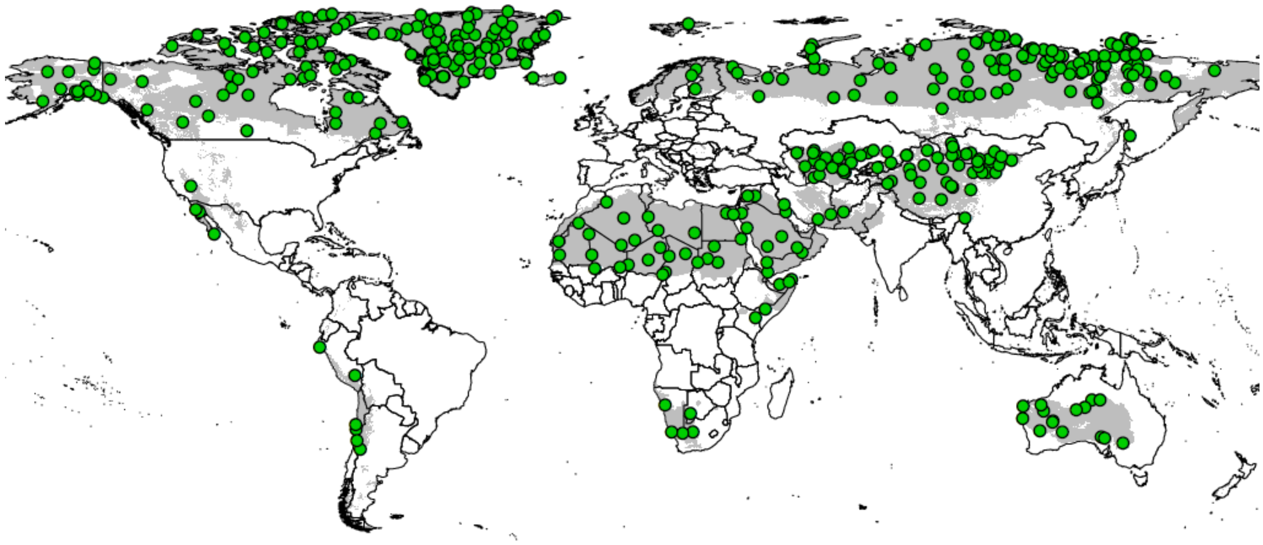

**Figure S14. Map showing locations of true absences (soybean yield equals zero) added to the historical yield dataset.** These data points (green dots) representing true absences of soybean were added to the dataset in order to extend the range of climate conditions captured by the model and to capture climate conditions leading to zero yield. These additional data points were randomly sampled in climate zones known to be unsuitable for soybean production (e.g. deserts and arctic areas) and added to the dataset with their yield value set to zero. Grey zones indicate climate zones from the Köppen-Geiger climate classification[21,22] in which the true absences were randomly selected. See Table S8 for a short description of these climate zones.

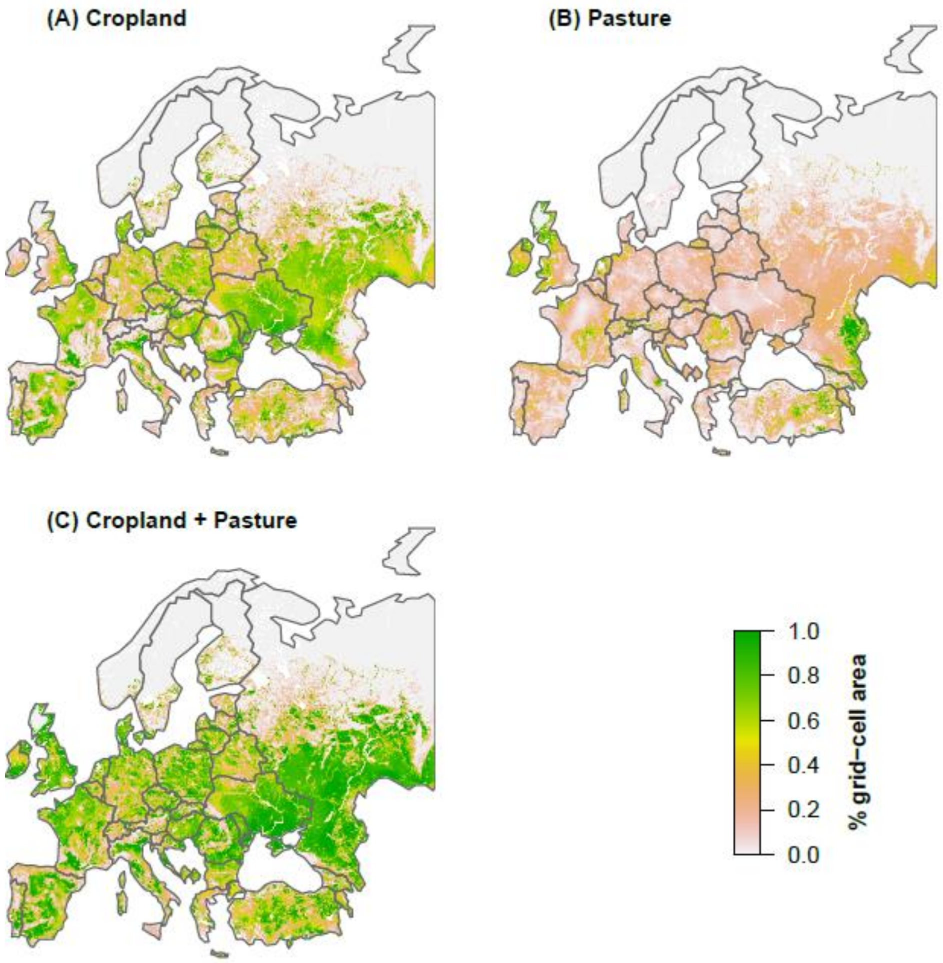

**Figure S15. Cropland (A), pasture (B), and total agricultural area (C) maps in Europe around the year 2000[23].** Total agricultural area is calculated as the sum of cropland plus pasture areas.

**Table S8. Climate zones from the Köppen-Geiger climate classification in which true absences (soybean yield equals zero) were randomly selected.** Locations of these climate zones can be seen in Figure S14.

| Climate zone code | Description of the corresponding climate |
|---|---|
| BWh | arid, winter dry , hot arid |
| BWk | arid, winter dry , cold arid |
| Dfc | snow, fully humid, cool summer |
| Dfd | snow, fully humid, extremely continental |
| EF | polar frost |
| ET | polar tundra |

### Supplementary details and discussion about the Random Forest model

#### *Analysis of residuals*

As shown in Figure S2, the predictive ability of the Random Forest algorithm is good. An analysis of residuals (yield data - yield prediction) reveals a tendency of the model to overestimate low yields and underestimate high yield (Figure S16-A), highlighting a conservative behavior of the model. Nevertheless, histogram of residuals indicates that the distribution of the residuals is symmetrical and centered, and that most residuals range between -1 to +1 t $ha^{-1}$ (Figure S16-B). Similar observations (good predictive ability and conservative behavior) about Random Forest when used to predict crop yields have been reported in the literature[24].

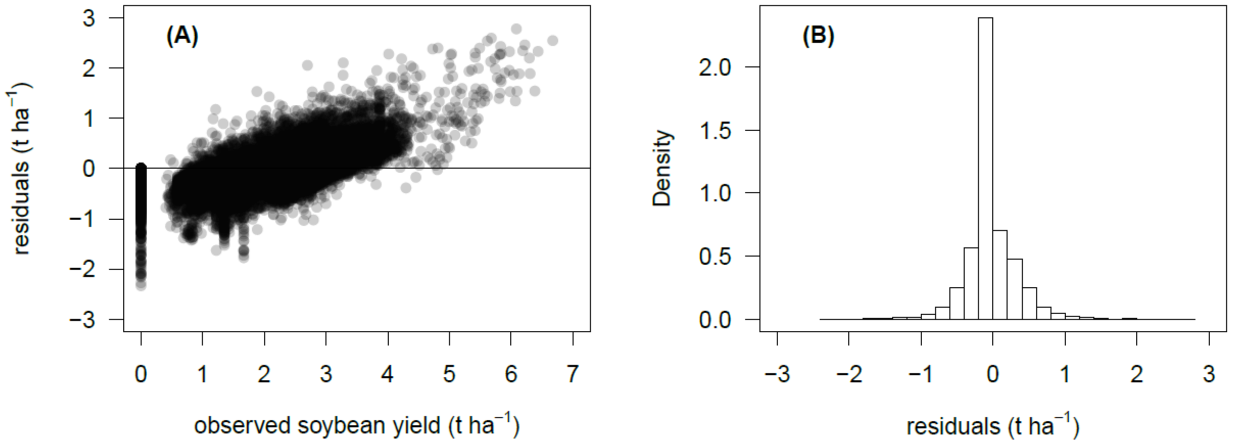

Figure S16. Analysis of the Random Forest residuals. (A) Residuals (data - prediction) as a function of observed soybean yield. (B) Histogram of residuals. Random Forest was fitted to the whole dataset.

### *Variable importance*

Variable importance of the fitted Random Forest algorithm is presented in Figure S17. It highlights the importance of temperature in the model, which is consistent with the key role of temperature in soybean suitable area shifts due to climate change found with the LDA (Figure 3).

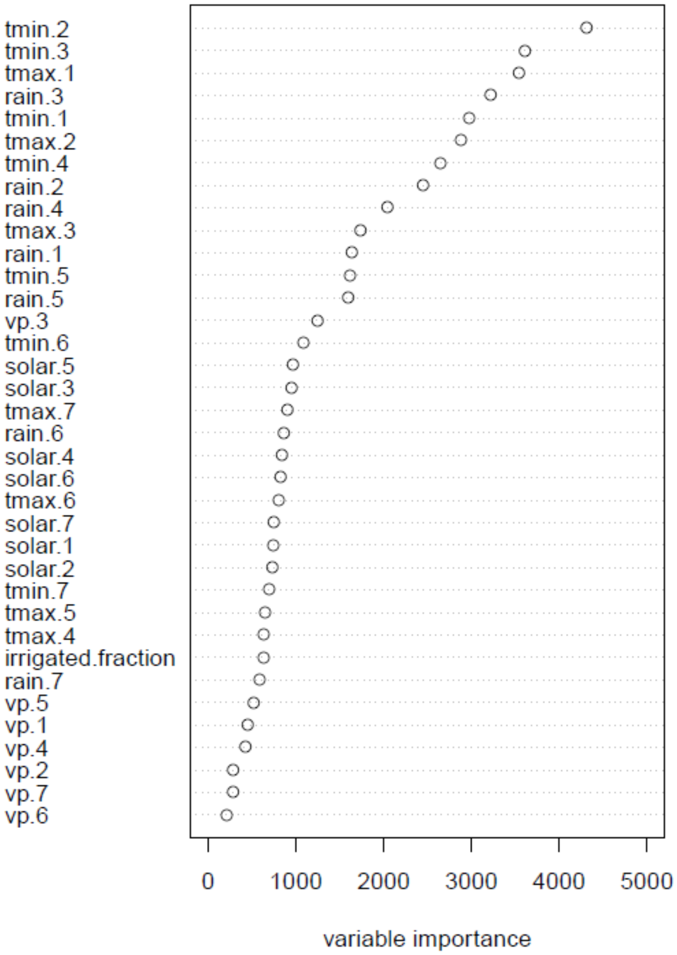

**Figure S17. Measures of importance of the inputs used by Random Forest .** The Random Forest algorithm (regression mode) was fitted with the function *ranger()* of the R package *Ranger* v0.10.1 with argument "importance" set to "impurity". Variable importance is assessed by a measure of impurity which is the variance of the responses. Variables are sorted by decreasing order. *tmax*: monthly average daily maximum air temperature at 2m, *tmin*: monthly average daily minimum air temperature at 2m, *rain*: monthly total precipitation (mm month$^{-1}$), *solar*: monthly average daily solar radiation (MJ m$^{-2}$ day$^{-1}$), *vp*: monthly average daily vapor pressure (hPa). The numerical suffix refers to the month of the soybean growing season (from 1 to 7).

### Additional references

1. Food and Agriculture Organization of the United Nations. FAOSTAT Statistics Database. (2019). Available at: http://www.fao.org/faostat/en/#data.
2. Iizumi, T., Okada, M. & Yokozawza, M. A meteorological forcing data set for global crop modeling: Development, evaluation, and intercomparison. *J. Geophys. Res. Atmos. Res.* **119**, 363–384 (2014).
3. Taylor, K. e., Stouffer, R. J. & Meehl, G. A. An Overview of CMIP5 and experiment design. *Am. Meteorol. Soc.* **93**, 485–498 (2012).
4. Monfreda, C., Ramankutty, N. & Foley, J. A. Farming the planet : 2. Geographic distribution of crop areas , yields , physiological types , and net primary production in the year 2000. *Global Biogeochem. Cycles* **22**, 1–19 (2008).
5. Mueller, N. D. *et al.* Closing yield gaps through nutrient and water management. *Nature* **490**, 254–257 (2012).
6. Kühling, I., Hüsing, B., Bome, N. & Trautz, D. Soybeans in high latitudes : effects of Bradyrhizobium inoculation in Northwest Germany and southern West Siberia. *Org. Agric.* **8**, 159–171 (2018).
7. Kadžiulienė, Ž., Arlauskienė, A. & Šarūnaitė, L. Legumes for sustainability of agroecosystems. in *20th Baltic Agronomy Forum* (eds. Gaile, Z. & Dace, Š.) 56 (2016).
8. Zimmer, S. *et al.* Effects of soybean variety and Bradyrhizobium strains on yield, protein content and biological nitrogen fixation under cool growing conditions in Germany. *Eur. J. Agron.* **72**, 38–46 (2016).
9. Pannecoucque, J. *et al.* Screening for soybean varieties suited to Belgian growing conditions based on maturity, yield components and resistance to Sclerotinia sclerotiorum and Rhizoctonia solani anastomosis group 2-2IIIB. *J. Agric. Sci.* 1–8 (2018). doi:10.1017/S0021859618000333
10. Lamichhane, J. R. *et al.* Analysis of soybean germination, emergence, and prediction of a possible northward establishment of the crop under climate change. *Eur. J. o* **113**, 125972 (2020).
11. Salem, M. A., Kakani, V. G., Koti, S. & Reddy, K. R. Pollen-Based Screening of Soybean Genotypes for High Temperatures. *Crop Sci.* **47**, 219–231 (2007).
12. Djanaguiraman, M., Schapaugh, W., Fritschi, F., Nguyen, H. & Prasad, P. V. Reproductive success of soybean (Glycine max L . Merril) cultivars and exotic lines under high daytime temperature. *Plant Cell Environ.* **42**, 321–336 (2019).
13. Harley, P. C., Weber, J. a & Gates, D. M. Interactive effects of light, leaf temperature, CO2 and O2 on photosynthesis in soybean. *Planta* **165**, 249–263 (1985).
14. Setiyono, T. D. *et al.* Simulation of soybean growth and yield in near-optimal growth conditions. *F. Crop. Res.* **119**, 161–174 (2010).
15. Hesketh, J. D., Myhre, D. L. & Willey, C. R. Temperature Control of Time Intervals Between Vegetative and Reproductive Events in Soybeans 1 . *Crop Sci.* **13**, 250–254 (1973).
16. Egli, D. B. & Wardlaw, I. F. Temperature Response of Seed Growth Characteristics of Soybeans. *Agron. J.* **72**, 560–564 (1980).

17. Boote, K. J. *et al.* Elevated temperature and CO2 impacts on pollination, reproductive growth, and yield of several globally important crops. *J. Agric. Meteorol.* **60**, 469–474 (2005).

18. Baker, J. T., Allen, L. H., Boote, K. J., Jones, P. & Jones, J. W. Response of Soybean to Air Temperature and Carbon Dioxide Concentration. *Crop Sci.* **29**, 98–105 (1989).

19. Brown, D. M. & Chapman, L. J. Soybean Ecology. II. Development-Temperature-Moisture Relationships from Field Studies. *Agron. J.* **52**, 496–499 (1960).

20. Iizumi, T. *et al.* Historical changes in global yields: Major cereal and legume crops from 1982 to 2006. *Glob. Ecol. Biogeogr.* **23**, 346–357 (2014).

21. Kottek, M., Grieser, C., Beck, C., Rudolf, B. & Rubel, F. World Map of the Köppen-Geiger climate classification updated. *Meteorol. Zeitschrift* **15**, 259–263 (2006).

22. Rubel, F., Brugger, K., Haslinger, K. & Auer, I. The climate of the European Alps: Shift of very high resolution Köppen-Geiger climate zones 1800-2100. *Meteorol. Zeitschrift* **26**, 115–125 (2017).

23. Ramankutty, N., Evan, A. T., Monfreda, C. & Foley, J. A. Farming the planet : 1. Geographic distribution of global agricultural lands in the year 2000. *Global Biogeochem. Cycles* **22**, 1–19 (2008).

24. Jeong, J. H. *et al.* Random Forests for Global and Regional Crop Yield Predictions. *PLoS One* **11**, e0156571 (2016).
